# A secreted NlpC/P60 endopeptidase from *Photobacterium damselae* subsp. *piscicida* cleaves the peptidoglycan of potentially competing bacteria

**DOI:** 10.1101/2020.06.30.181511

**Authors:** Johnny Lisboa, Cassilda Pereira, Aline Rifflet, Juan Ayala, Mateus S. Terceti, Alba V. Barca, Inês Rodrigues, Pedro J.B. Pereira, Carlos R. Osorio, Francisco García-del Portillo, Ivo Gomperts Boneca, Ana do Vale, Nuno M.S. dos Santos

**Affiliations:** Fish Immunology and Vaccinology Group, IBMC – Instituto de Biologia Molecular e Celular, Universidade do Porto, Porto, Portugal; Fish Immunology and Vaccinology Group, i3S - Instituto de Investigação e Inovação em Saúde, Universidade do Porto, Porto, Portugal; Institut Pasteur, Unité Biologie et Génétique de la Paroi Bactérienne, Paris, France; INSERM Groupe Avenir, Paris, France; CNRS, UMR “Integrated and Molecular Microbiology”, Paris, France; Centro de Biología Molecular Severo Ochoa, Consejo Superior de Investigaciones Científicas (CBMSO-CSIC), Madrid, Spain; Departamento de Microbioloxía e Parasitoloxía, Instituto de Acuicultura, Universidade de Santiago de Compostela, Santiago de Compostela, Spain; Biomolecular Structure Group, IBMC – Instituto de Biologia Molecular e Celular, Universidade do Porto, Portugal; Macromolecular Structure Group, i3S - Instituto de Investigação e Inovação em Saúde, Universidade do Porto, Porto, Portugal; Laboratorio de Patógenos Bacterianos Intracelulares, Centro Nacional de Biotecnología, Consejo Superior de Investigaciones Científicas (CNB-CSIC), Madrid, Spain

**Author notes:** Address correspondence to Nuno M.S. dos Santos, or Johnny Lisboa. Cassilda Pereira, Stem Cells in Regenerative Biology and Repair, INEB - Instituto Nacional de Engenharia Biomédica, Universidade do Porto, Portugal and i3S - Instituto de Investigação e Inovação em Saúde, Universidade do Porto, Porto, Portugal. Mateus S. Terceti, Departamento de Biologia Geral e Aplicada, Instituto de Biociências de Rio Claro – São Paulo, Universidade Estadual Paulista – ENESP, Brasil.

## Abstract

**Peptidoglycan** (PG) is a major component of the bacterial cell wall, forming a mesh-like structure enwrapping the bacteria that is essential for maintaining structural integrity and providing support for anchoring other components of the cell envelope. PG biogenesis is highly dynamic and requires multiple enzymes, including several hydrolases that cleave glycosidic or amide bonds in the PG. Here, it is described the structural and functional characterization of an NlpC/P60-containing peptidase from *Photobacterium damselae* subsp. *piscicida* (*Phdp*), a Gram-negative bacterium that causes high mortality of warm-water marine fish with great impact for the aquaculture industry. PnpA (*<u>P</u>hotobacterium* <u>N</u>lpC-like <u>P</u>rotein <u>A</u>) has a four-domain structure with a hydrophobic and narrow access to the catalytic center and specificity for the γ-D-glutamyl-meso-diaminopimelic acid bond. However, PnpA does not cleave the PG of *Phdp* and neither PG of several Gram-negative and Gram-positive bacterial species. Interestingly, it is secreted by the *Phdp* type II secretion system and degrades the PG of *Vibrio anguillarum* and *V. vulnificus*. This suggests that PnpA is used by *Phdp* to gain an advantage over bacteria that compete for the same resources or to obtain nutrients in nutrient-scarce environments. Comparison of the muropeptide composition of PG susceptible and resistant to the catalytic activity of PnpA, showed that the global content of muropeptides is similar, suggesting that susceptibility to PnpA is determined by the three-dimensional organization of the muropeptides in the PG.

**IMPORTANCE:** Peptidoglycan (PG) is a major component of the bacterial cell wall formed by long chains of two alternating sugars interconnected by short peptides, originating a mesh-like structure that enwraps the bacterial cell. Although PG provides structural integrity and support for anchoring other components of the cell envelope, it is constantly being remodeled through the action of specific enzymes that cleave or joint its components. Here, it is shown that *Photobacterium damselae* subsp. *piscicida*, a bacterium that causes high mortality in warm-water marine fish, produces PnpA, an enzyme that is secreted into the environment and is able to cleave the PG of potentially competing bacteria, either for gaining competitive advantages and/or to get nutrients. The specificity of PnpA to the PG of some bacteria and its inability to cleave others may be explained by differences in the structure of the PG mesh and not by different muropeptide composition.

## INTRODUCTION

Peptidoglycan (PG) is a major component of the bacterial cell wall, essential for maintaining structural integrity and internal osmotic pressure, shaping the morphology of bacteria and providing support for anchoring other components of the cell envelope (1, 2). PG forms a mesh-like structure that enwraps the bacterial cell, referred to as sacculus, which is composed of long chains of two alternating β(1-4) glycosidic-bonded glycans, N-acetylglucosamine (GlcNAc) and N-acetylmuramic acid (MurNAc), cross-linked by short stem peptides, either directly or through bridging peptides (1, 3-5). The stem peptides are usually 4-5 amino acids long, contain L- and D-amino acids and extend from MurNAc (1-4). The most common structure of the stem peptide is L-Ala-γ-D-Glu-m-DAP-D-Ala-D-Ala (m-DAP standing for *meso*-diaminopimelic acid) in Gram-negative bacteria and L-Ala-γ-D-Glu-L-Lys-D-Ala-D-Ala] in Gram-positive organisms (1, 2, 4).

In spite of its stabilizing function, PG is highly dynamic, with covalent bonds being formed and broken by different enzymes. Multiple hydrolases, capable of cleaving glycosidic (glycosidases) or amide (amidases and peptidases) bonds in the PG sacculus and/or its soluble fragments, play a preponderant role in PG dynamics (1, 2, 6-13). Degradation products resulting from the catalytic activity of PG hydrolases can be recycled for PG *de novo* biosynthesis and also act as signaling molecules in quorum sensing, triggering antibiotic resistance, regrowth of dormant cells or as effector molecules in immune responses (1, 2, 6, 7, 12, 14, 15). Besides their role in PG dynamics, hydrolases can also be secreted to the environment or injected via type VI secretion systems into the periplasm of other bacteria to confer competitive advantage over competing bacteria that share mixed growth environments or as a way of obtaining nutrients (1, 10, 11, 16-23).

PG peptidases are a widely diverse group of enzymes, with ten different types of catalytic domains involved in PG hydrolysis described to date (1, 24). Amongst them, cysteine peptidases containing New lipoprotein C/Protein of 60-kDa (NlpC/P60) catalytic domains are present in most bacterial lineages, suggesting that they play an important biological role (1, 24). NlpC/P60-containing peptidases are involved in the catalysis of the N-acetylmuramate-L-alanine or D-γ-glutamyl-mesodiaminopimelate linkages, with four major groups identified so far: (i) P60-like, (ii) AcmB/LytN-like, (iii) YaeF/Poxvirus G6R, and (iv) lecithin retinol acyltransferase (LRAT)-like (24). The NlpC/P60 domain is structurally similar to a primitive papain-like peptidase (24-29) and can be found alone or fused to other domains, with or without catalytic functions, to form multifunctional proteins (1, 2, 24, 26, 30-35). Several of these domains, such as the SH3 (sarcoma homology 3) domain (31, 32, 35), are involved in anchoring hydrolases to cell wall components, allowing their appropriate concentration and positioning for the formation of an efficient enzyme-substrate complex (1).

*Photobacterium damselae* subsp. *piscicida* (*Phdp*) is a gram-negative, halophilic bacterium that induces an acute infection with development of a rapid septicemia, resulting in high mortality of warm-water marine fish with devastating consequences for the aquaculture industry (36, 37). Although it has been suggested that *Phdp* remains in a cultivable form in salt water for only 4-5 days (38, 39), it was also suggested that it has the ability to enter a dormant, non-cultivable but infectious state in salt water and sediment (40). With regard to the mechanisms responsible for the pathogenicity of *Phdp*, it was shown that extracellular products (ECPs) play a fundamental role (41, 42) although among their components, only the toxin AIP56 has been identified and characterized so far (43-47).

The present work reports the structural and functional characterization of a novel NlpC/P60-containing peptidase from *Phdp* (PnpA). The results show that PnpA is a PG hydrolase with a four-domain structure similar to that of DvLysin and specificity for the γ-D-glutamyl-meso-diaminopimelic acid bond (26), but with a more hydrophobic and narrower access to the catalytic center. It is also shown that PnpA is secreted into the extracellular medium by the *Phdp* type II secretion system and acts on the PG of *Vibrio anguillarum* and *V. vulnificus*, suggesting that it may provide to *Phdp* an advantage over bacteria competing for the same resources or a way of obtaining nutrients in nutrient-scarce environments, either inside or outside the host. Comparison of the muropeptide compositions of the PG, susceptible and resistant to PnpA activity, allowed to develop a model suggesting that the susceptibility to PnpA is determined by three-dimensional structural features of the PG and not by their chemical compositions.

## RESULTS

### *Photobacterium damselae* subsp. *piscicida* (*Phdp*) secretes an NlpC/P60 family protein

*Phdp* virulent strains have a relatively simple profile of secreted proteins in mid-exponential growth phase cultures (45). Apart from AIP56 toxin, no other proteins have been identified and characterized. In this work, a protein band of approximately 55 kDa was excised from an SDS-PAGE gel and subjected to MALDI-TOF-MS. The obtained MS data were used in a Mascot search against the NCBI database resulting in the identification of a hypothetical protein from *Photobacterium damselae* subsp. *damselae* (*Phdd*) CIP 102761 (VDA_000779; accession: EEZ39759). The 1479 nucleotide homologous sequence in the *Phdp* MT1415 strain (accession: TJZ86030.1) was then amplified using primers designed based on the VDA_000779 sequence. *In silico* analysis (SignalP 5.0 and NCBI Conserved domain search) of its 499 amino-acids translation product predicted a Sec signal peptide (M^1^ to A^19^), followed by an N_NLPC_P60 putative stabilizing domain (Pfam PF12912), an SH3b1(Pfam PF12913/12914) and an NlpC_P60 domain (Pfam PF00877), classifying it as a protein belonging to the NlpC/P60 family, hereinafter referred to as PnpA (*<u>P</u>hotobacterium* <u>N</u>lpC-like <u>P</u>rotein <u>A</u>).

### PnpA is encoded in a genetically unstable chromosomal region and its expression levels are similar at exponential and stationary phases of growth

To investigate the genetic context of *pnpA* in *Phdp* MT1415 strain, the draft genome sequence of MT1415 was obtained in this study. Then, homologous DNA sequences of a number of *Phdp* and *Phdd* isolates were additionally retrieved from the GenBank database and subjected to comparative sequence analysis (Fig. 1A). This evidenced that the PnpA-encoding gene is invariably linked to a downstream gene encoding a Ribonuclease T, and to an upstream gene encoding an α-galactosidase, the latter being a pseudogene in some *Phdp* isolates. As a whole, the DNA flanking *pnpA* underwent a massive insertion of transposase genes (IS elements of the IS*1* and IS*91* families) likely followed by accumulation of inactivating mutations that rendered a collection of pseudogenes. This process of gene decay not only affected the transposase genes themselves, but also flanking genes encoding enzymes putatively involved in sugar metabolism, as α-galactosidases, α-amylases and pullulanases (Fig. 1A). Proliferation of insertion sequences that cause high frequency of pseudogenes and gene loss is indeed a hallmark of all *Phdp* genomes studied to date (48-50). The observation that PnpA and the Ribonuclease T genes have escaped the inactivation by IS insertions, suggest that these two genes may fulfill an important role in *Phdp*.

**FIG 1.**
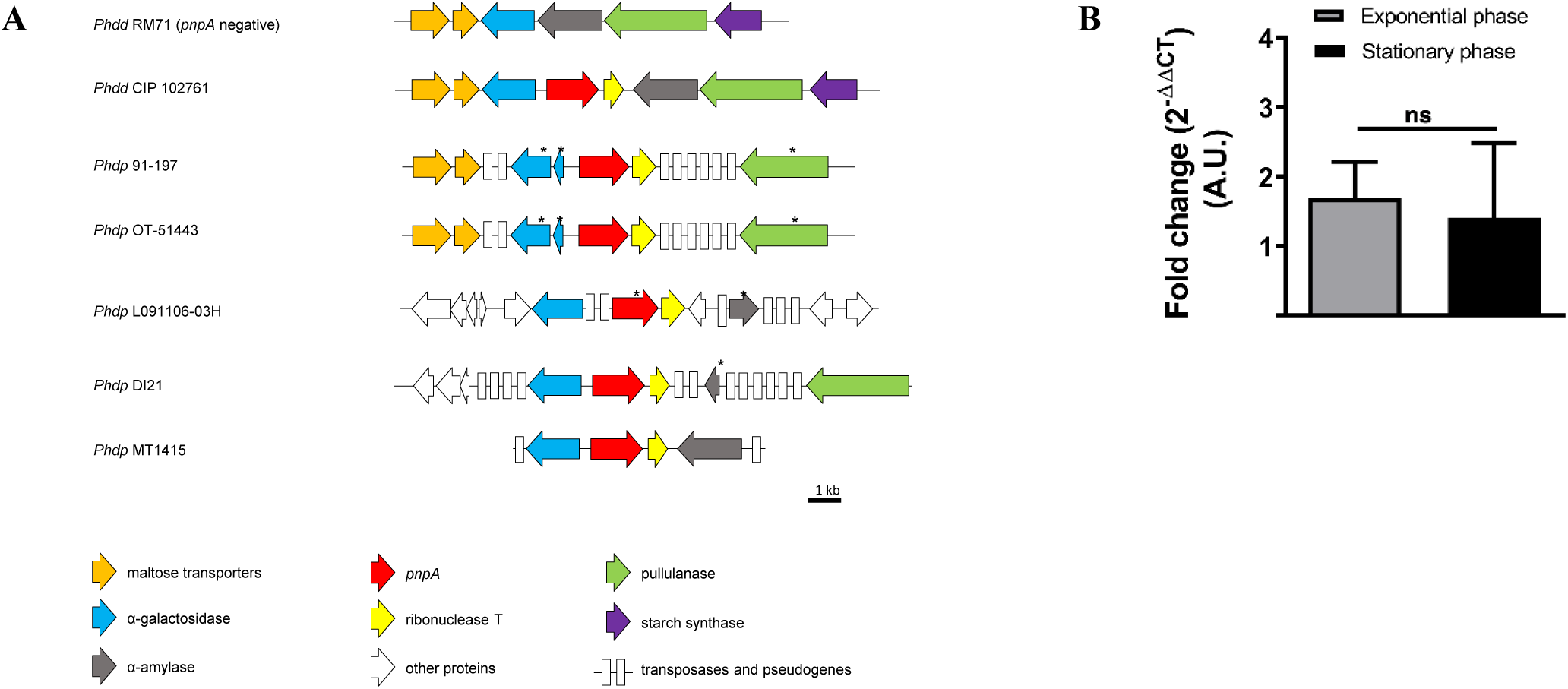
Genomic context and expression of pnpA. (A) Schematic representation of the genomic context of pnpA (shown in red) in the indicated Photobacterium damselae subsp. damselae (Phdd) and Photobacterium damselae subsp. piscicida (Phdp) strains. * Denotes truncated gene version. (B) pnpA expression in vitro. Expression levels in Photobacterium damselae subsp. piscicida MT1415 strain at exponential (OD_600_ of 0.4) and stationary (OD_600_ of 1.2) phases were determined using quantitative RT-PCR, normalized based on expression of the housekeeping gene 16S and expressed as 2-ΔΔCT. Data represent the mean ± SD of three independent experiments with all measurements done in triplicate. Statistical significance between expression at exponential and stationary phase was assessed using Student’s t-test. ns = non-significant.

Expression levels of *pnpA* were determined by RT-PCR, showing that under the culture conditions used (growth in TSB-1 at 25°C), there are no differences in the level of gene transcription between exponential and stationary phase cultures (Fig. 1B).

### Overall description of PnpA structure

For better understanding the structure-function of PnpA, its three-dimensional structure was solved. The crystal structure of PnpA was determined at 1.4 Å resolution by molecular replacement with DvLysin (PDB entry 3M1U, 26% sequence identity), an endopeptidase from *Desulfovibrio vulgaris* Hildenborough (26). The asymmetric unit contains two PnpA molecules, which are essentially identical (root-mean-square-deviation [r.m.s.d], of 0.5 Å for 457 aligned Cα atoms). Table S1 summarizes the data collection, processing, and refinement statistics.

Analysis of the intermolecular packing interfaces within the crystal lattice suggest that the molecule behaves as a monomer in solution, which is in agreement with the molecular mass estimated by size-exclusion chromatography. The PnpA monomer has an overall structure similar to that of DvLysin (26), namely, one N-terminal “c-clip” or “N_NLPC_P60” stabilizing domain (residues N^20^-N^133^), two SH3b domains (SH3b1, residues I^134^-V^218^; SH3b2, residues D^219^-T^295^), and the C-terminal NlpC/p60 catalytic domain (residues P^296^-K^499^) (Fig. 2A). The three-dimensional models of DvLysin and PnpA display an r.m.s.d of 2.2 Å (for 405 aligned Cα atoms), suggesting that both proteins may be functionally equivalent. A significant number of structures sharing at least one of the PnpA domains have been identified (Table S2), although so far, PnpA and DvLysin are the only four-domain NlpC/P60-containing peptidases whose structure has been reported.

**FIG 2.**
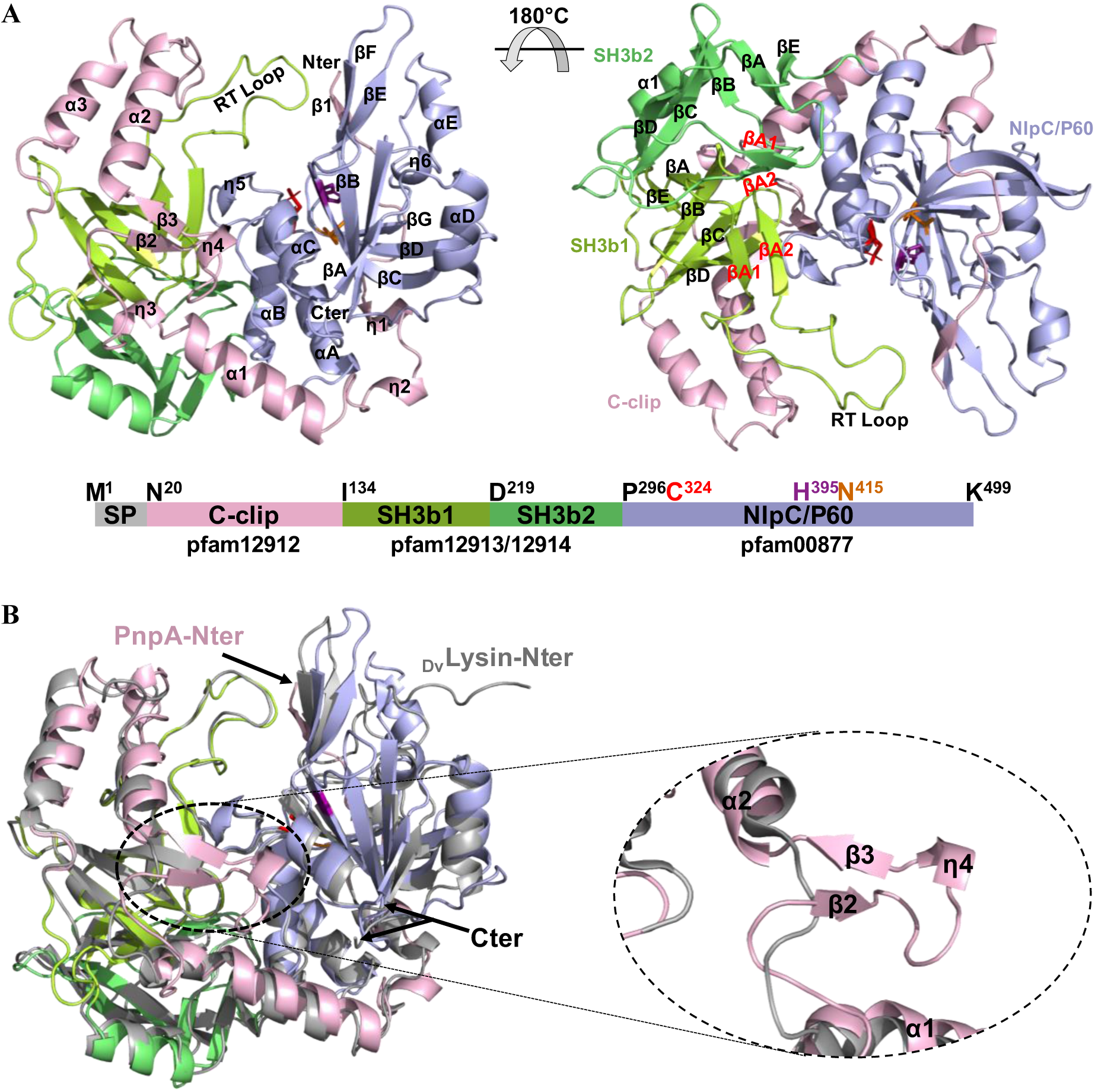
Three-dimensional structure of Phdp PnpA. (A) Cartoon representation of the PnpA monomer, with domains colored as in the linear representation shown below (signal peptide (SP) gray; C-clip domain pink; SH3b1 domain light green; SH3b2 domain dark green; NlpC/P60 domain blue; domain boundaries and catalytic residues are indicated). The catalytic site residues are represented as sticks (C324 red, H395 pink and N415 orange). N and C termini and secondary structure elements are labeled. (B) Cartoon representation of superposed PnpA (color code as in panel A) and DvLysin (grey). N- and C-termini are indicated. A close-up of the insertion between α1 and α2, forming an additional antiparallel β-sheet (β2 and β3) and a 310 helix (η4) in the c-clip domain, is shown in the insert (dashed oval).

As in DvLysin (26), the PnpA c-clip domain has an extended helical conformation which surrounds and stabilizes the SH3b1 and NlpC/P60 domains, forming a planar assembly from which the SH3b2 domain protrudes (Fig. S1). Compared to DvLysin, the c-clip domain of PnpA harbors an extension between helices α1 and α2, thereby forming an additional two-stranded antiparallel β-sheet (β2 and β3) and a 3_10_ helix (η4), which protrude into the catalytic groove and close one of its sides (Fig. 2B).

The presence of SH3b domains in prokaryotes has long been documented. These domains have been described as targeting domains, involved in cell-wall recognition and binding (1, 24, 35). Despite the low amino acid sequence conservation (8% sequence identity), the two SH3b domains in PnpA have a conserved overall fold (r.m.s.d. of 3.9 Å for 55 aligned Cα atoms) (Fig. S2). As in DvLysin (26), both PnpA SH3b domains consist of seven conserved strands (βA-βA1-βA2-βB-βC-βD-βE), with the βA-βE strands structurally equivalent to their eukaryotic counterparts (31, 32), while the βA1 and βA2 form a β-hairpin that corresponds to the RT loops of eukaryotic SH3b domains (Fig. 2A).

As in other NlpC/P60-containing peptidases, the 204 residue-long C-terminal NlpC/P60 catalytic domain of PnpA displays a fold resembling a primitive papain-like cysteine peptidase (24). Its secondary structure elements adopt the topology described for DvLysin, i.e., a six-stranded central β sheet and five α helices with αA-αB-αC-βA-αD-βB-βC-βD-βE-αE-βF topology, where αA-αB-αC and αD-αE protect either side of the central β-sheet (Fig. 2A) (26).

### PnpA has a narrow and hydrophobic access to the catalytic site

The active site of NlpC/P60 cysteine peptidases consists of a conserved cysteine-histidine dyad and a third polar residue (H, N or Q) that orients and polarizes the catalytic histidine (24-29). In PnpA, the residues that make up the active site are C324, H395 and N415, the latter similar to the equivalent residue found in the active site of the prototypical papain (51), but differing from the histidine (H408) at the active site of DvLysin’s (26) (Fig. 3A). As described for other NlpC/P60-containing peptidases (24-29), the catalytic C324 is located at the amino terminus of a helix packing against the central β-sheet that harbors H395 in its second strand and N415 in the third. In the PnpA structure, the thiol group of the catalytic cysteine is oxidized, resulting in the disruption of the characteristic C324 SD-H395 ND1 hydrogen bond and suggesting that the enzyme is in an inactive state (Fig. S3). As advanced for BtYkfC (26), oxidation of the catalytic cysteine most likely occurred during crystallization or exposure to X-rays (52), since recombinant PnpA from the same purification batch was used in biochemical assays and was catalytically active.

**FIG 3.**
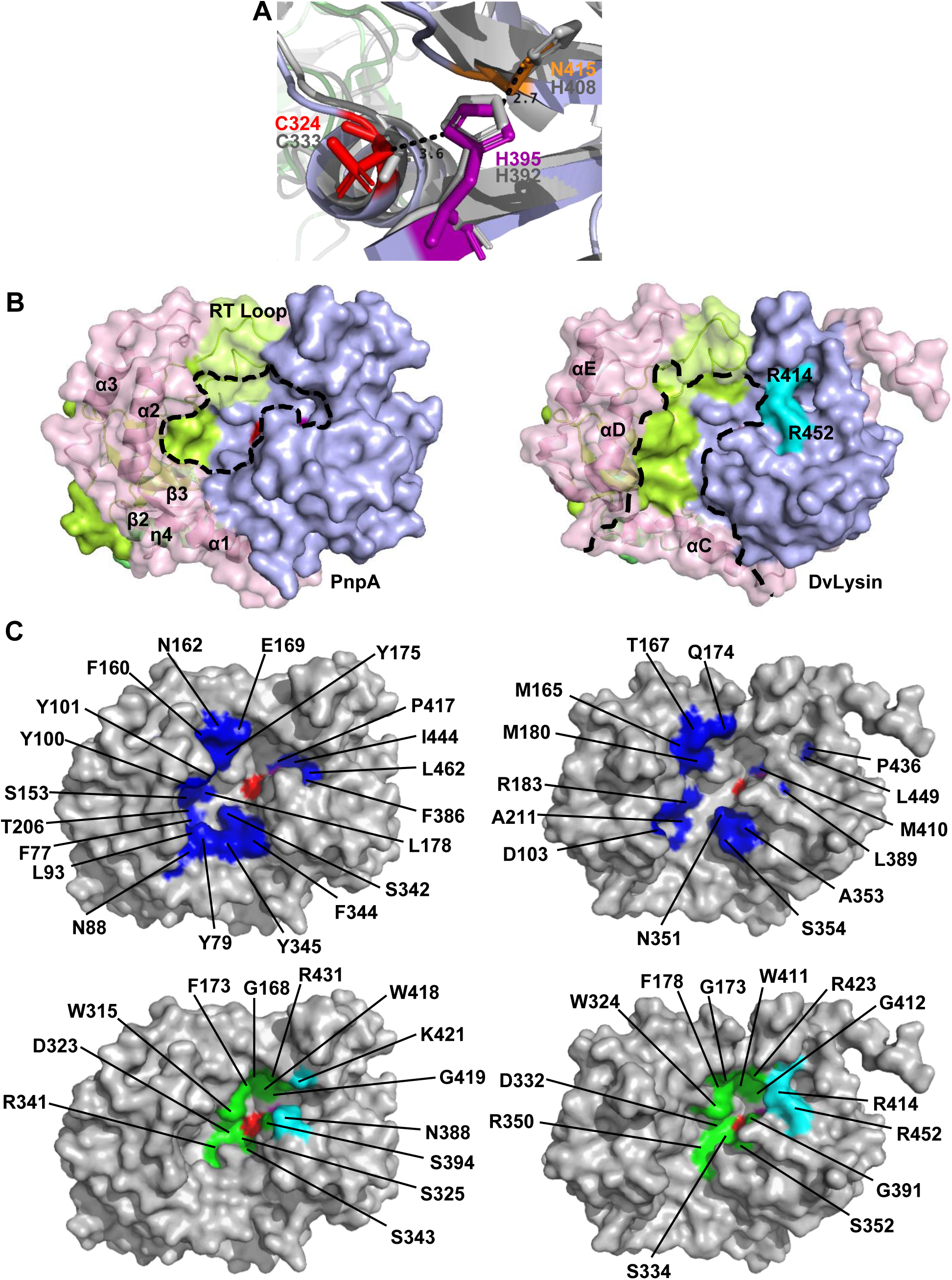
Structural comparison of the active sites of PnpA and DvLysin. (A) Superposition of the catalytic site of PnpA (colored sticks) and DvLysin (grey sticks). (B) Solid surface representation of PnpA (left) and DvLysin (right). Catalytic grooves are outlined by dashed lines. Residues R414 and R452 from DvLysin are colored cyan and labeled. (C) Comparison of the catalytic cavities of PnpA (left) and DvLysin (right). Hydrophobic and polar residues close to the substrate binding region are colored dark blue (top panel). DvLysin residues involved in substrate binding (27) and conserved in PnpA are colored green (lower panel). Catalytic residues are colored as in Fig. 2A.

In DvLysin, access to the catalytic cysteine occurs through a groove between the NlpC/P60 domain on one side and the c-clip helices αD and αE plus the SH3b1 domain on the other, with the RT loop from the SH3b1 domain closing one end of the groove (Fig. 3B) (26). While this topology is generally maintained in PnpA, the end of the groove opposite to the RT loop is also closed by strands β2 and β3 and the 3_10_ helix η4, creating a narrower access to the catalytic site (Fig. 3B). A minor difference is observed on the “wall” formed by the NlpC/p60 domain, wider in PnpA and closed by R414 and R452 in DvLysin. Besides the narrower entrance, two clusters of amino acids confer to the active site cavity of PnpA a more polar and hydrophobic nature than observed for DvLysin (Fig. 3C). However, extensive conservation of substrate-interacting residues between PnpA and DvLysin (Fig. 3C) suggest a similar interaction with mDAP-D-Ala from the stem peptide.

### PnpA is secreted by *Phdp* type II secretion system (T2SS)

PnpA possesses a typical Sec signal peptide and was identified in the culture supernatants of exponentially growing *Phdp* cultures, suggesting that it could be actively secreted by the bacteria. Many proteins that are transported via the Sec system into the periplasm are secreted across the outer membrane through a T2SS (53, 54). Recently, it was shown that *Phdp* contains a functional T2SS (44) and that deletion of *epsL*, which encodes an inner membrane-spanning protein that establishes a critical link between the cytoplasmic and the periplasmic parts of that system (55), abolishes the secretion of AIP56 (44). To test the involvement of the T2SS of *Phdp* in PnpA secretion, the presence of PnpA in total cell lysates and extracellular products of WT, Δ*epsL* and Δ*epsL +* pEpsL *Phdp* was analyzed by Western blotting (Fig. 4). PnpA was detected in ECPs, but not in total cell lysates of the WT strain, confirming that it is a secreted protein (Fig. 4A). In contrast, in the Δ*epsL* strain, PnpA was retained in the periplasm (Fig. 4B), confirming the involvement of T2SS in PnpA secretion.

**FIG 4.**
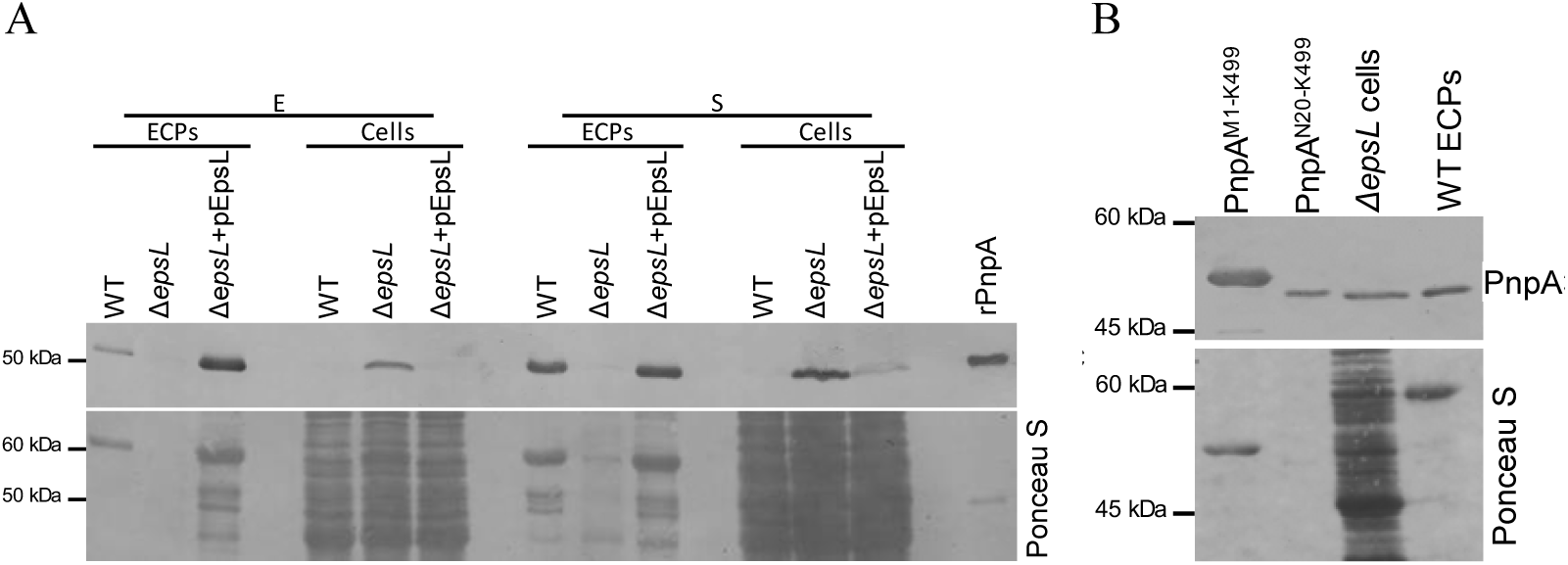
Secretion of PnpA is dependent on the type II secretion system (T2SS). (A) Wild type, ΔepsL and ΔepsL complemented (ΔepsL + pEpsL) strains grown to an OD_600_ of 0.5 (exponential phase, E) or 1.5 (stationary phase, S). Extracellular products (ECPs) and bacterial pellets (Cells) were subjected to SDS-PAGE and PnpA was detected by western blotting (upper panel). Recombinant PnpA (rPnpA; 0.2 µg) was used as a positive control. Lower panel, total protein loading (Ponceau S). Blot shown is representative of three independent experiments. (B) In ΔepsL cells, PnpA is retained at the periplasm. Western blotting analysis of PnpA retained in ΔepsL cells and secreted by the WT bacteria (upper panel). rPnpA containing and lacking the Sec signal peptide (PnpAM1-K499 and PnpAN20-K499, respectively) were run as references. Please note that PnpA retained in ΔepsL cells migrates similarly to the PnpA secreted by the WT bacteria, confirming the removal of the signal peptide and, thus, the periplasmic localization of PnpA in ΔepsL cells. Lower panel, total protein loading (Ponceau S).

### PnpA has specificity for the γ-D-glutamyl-meso-diaminopimelic acid bond

To investigate the PnpA enzymatic activity towards PG muropeptides and define its substrate specificity, recombinant PnpA was incubated with monomeric tri-(M3: GlcNAcMurNAc-L-Ala-D-Glu-mDap), tetra-(M4: GlcNAc-MurNAc-L-Ala-D-Glu-mDap-D-Ala) and penta-muropeptides (M5: GlcNAc-MurNAc-L-Ala-D-Glu-mDap-D-Ala-D-Ala) and the cleavage product(s) analysed by HPLC (Fig. 5). PnpA converted all tested muropeptides to dipeptides (M2: GlcNAc-MurNAc-L-Ala-D-Glu), suggesting it cleaves specifically γ-D-glutamyl-*meso*-diaminopimelic acid bond of monomeric muropeptides.

**FIG 5.**
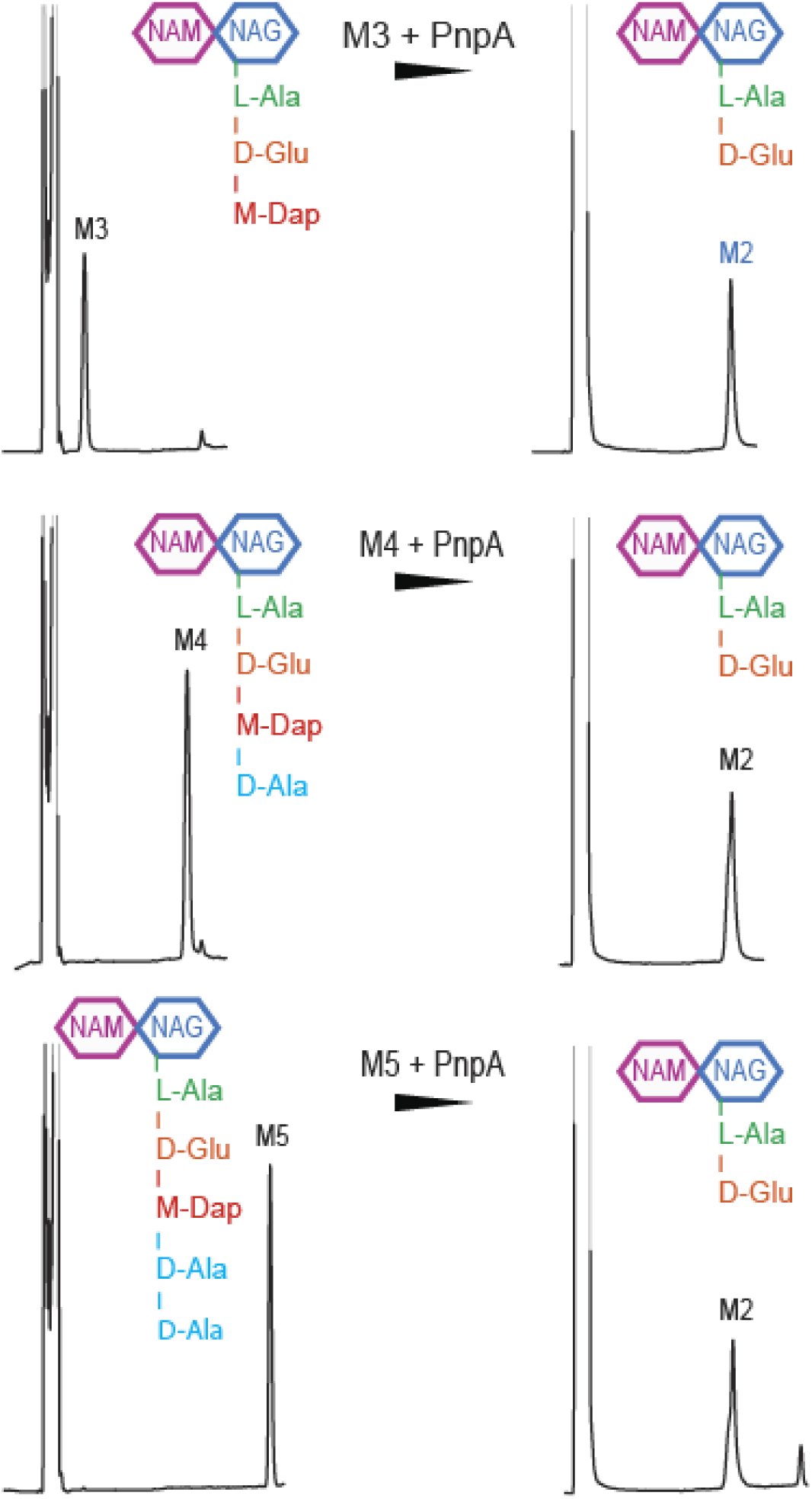
PnpA cleaves monomeric muropeptides M3, M4 and M5. HPLC profiles of each muropeptide at time 0 (left) and after 3 h incubation (right) with 50 µg mL^-1^ recombinant PnpA (rPnpA). Note that after incubation with PnpA, M2 was always obtained as a product. M3: GlcNAc-MurNAc-L-Ala-D-Glu-dap-D-Ala-D-Ala; M4: GlcNAc-MurNAc-L-Ala-D-Glu-dap-D-Ala; M5: GlcNAc-MurNAc-L-Ala-D-Glu-dap-D-Ala-D-Ala.

### PnpA does not hydrolyse *Phdp* peptidoglycan

In order to evaluate the involvement of PnpA in *Phdp* cell wall biogenesis, a *Phdp* Δ*pnpA* strain was generated (Figs. 6A-B). Bacterial growth was not affected in the deleted strain (Fig. 6C). In addition, no differences were detected in the composition of the peptidoglycan from the WT and Δ*pnpA* strains (Fig. 6D; Table 1). In agreement with this, both WT and Δ*pnpA* strains showed similar morphology (Fig. 6E). Moreover, PnpA did not display *in vitro* enzymatic activity against *Phdp* whole sacculus, since no differences in the muropeptide composition were detected after incubating the PG with active PnpA or inactive PnpA (Figs. 6F, S4A and S5A; Table 2). Altogether, these results suggest that PnpA is not enzymatically active towards *Phdp* PG.

**TABLE 1.**
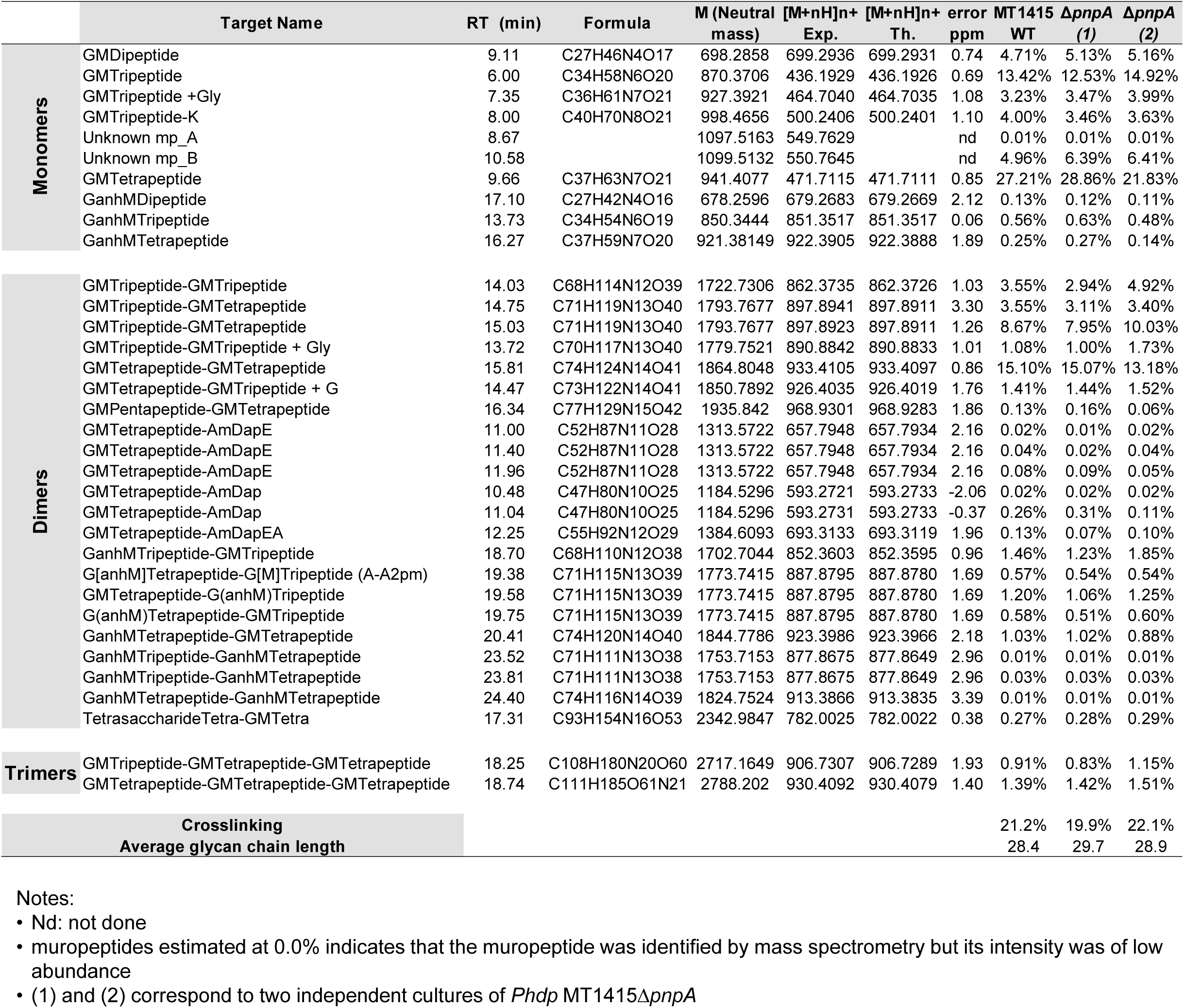
Structure, molecular mass and quantity of muropeptides from wild type and Δ*pnpA Phdp* MT1415 strains

**TABLE 2.**
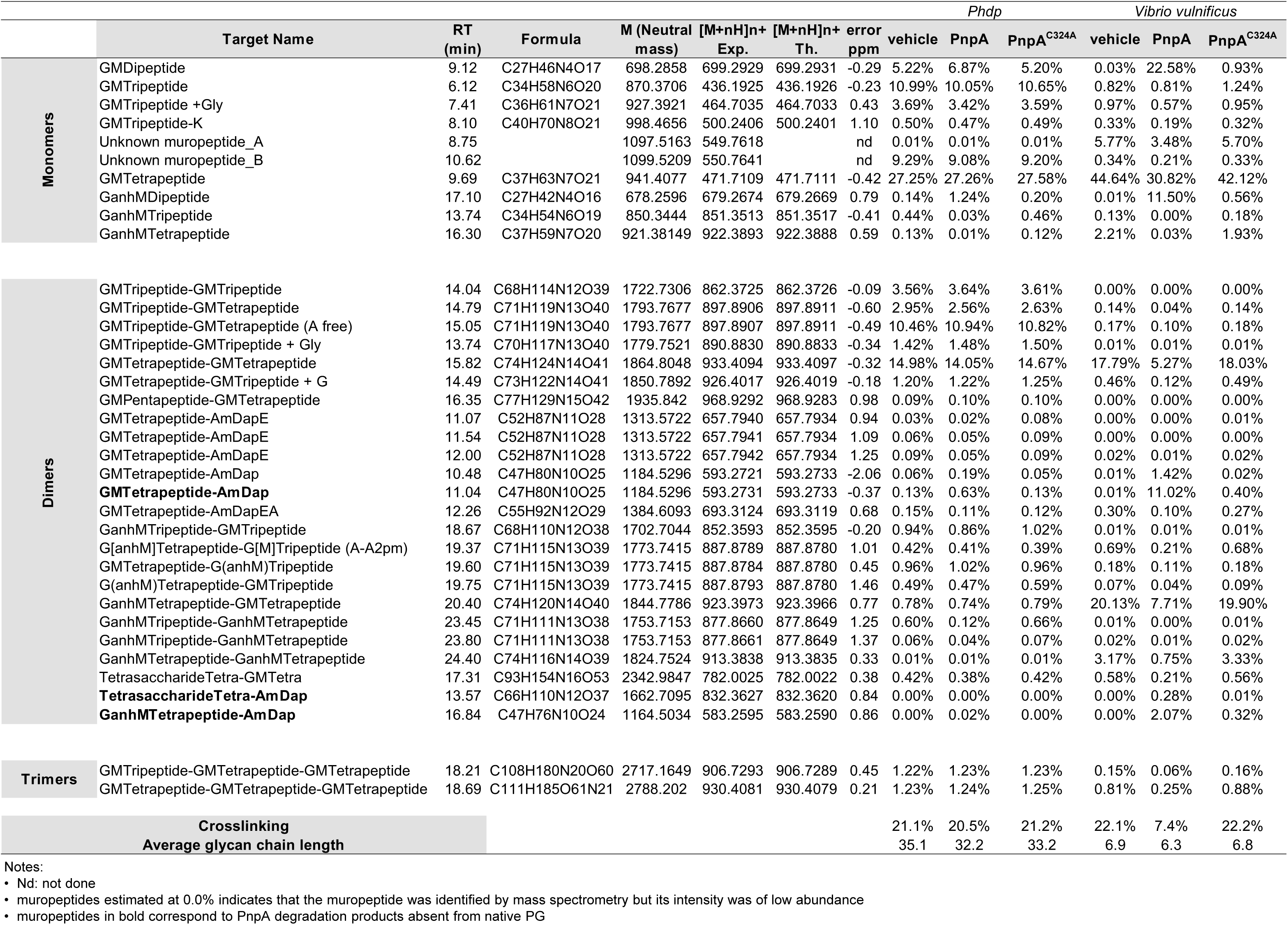
Structure, molecular mass and quantity of muropeptides from *Phdp* and *V. vulnificus* PG incubated with vehicle, PnpA or PnpA^C324A^

**FIG 6.**
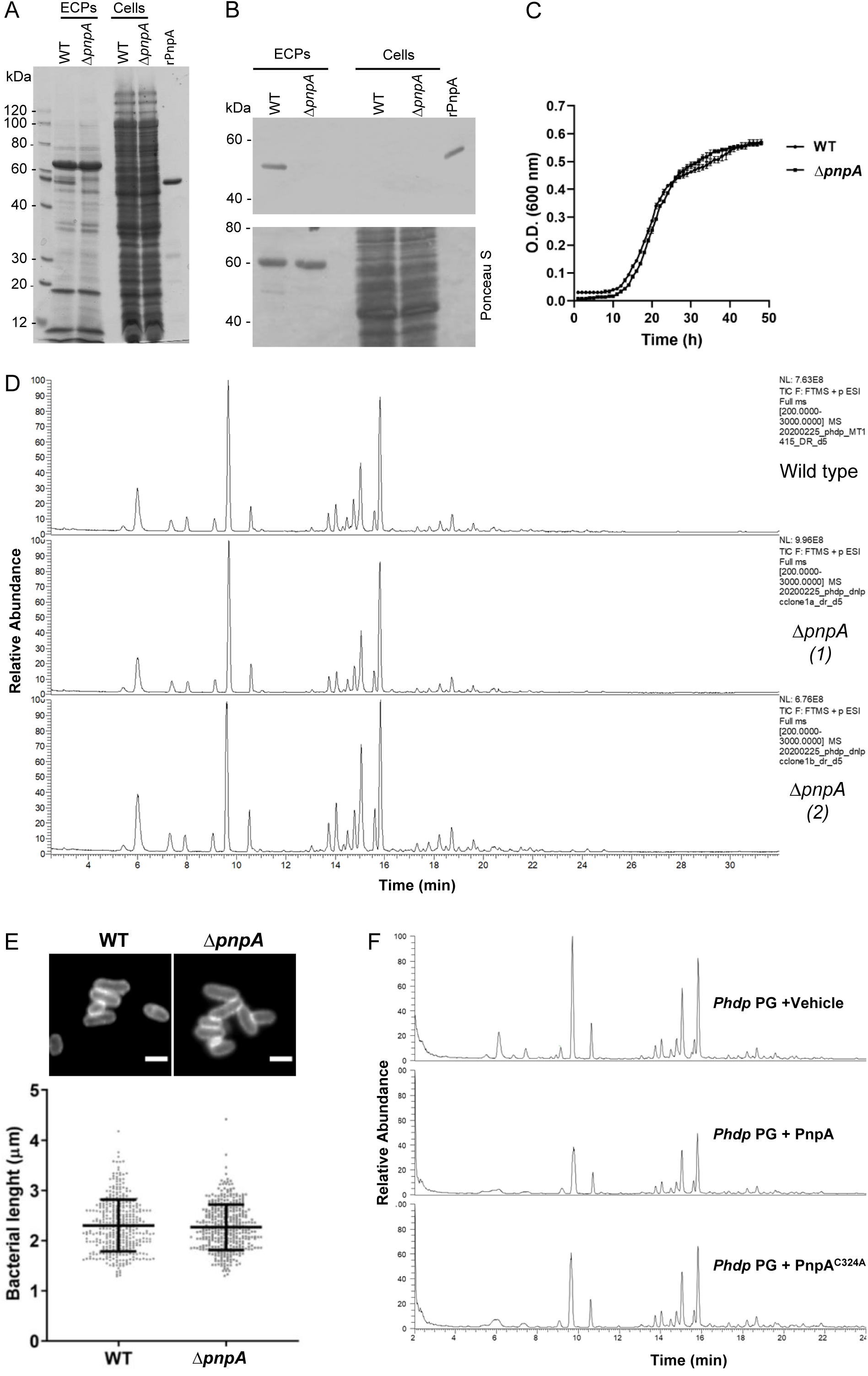
PnpA does not hydrolyses Phdp peptidoglycan. (A) SDS-PAGE of extracellular products (ECPs) and bacterial pellets (Cells) from WT and ΔpnpA Phdp. ECPs equivalents to 1.5 mL and cells equivalent to 0.3 mL of early stationary culture were separated by 12% SDS-PAGE and stained with Coomassie. Recombinant PnpA (rPnpA; 2 µg) was used as reference. Gel shown is representative of two independent experiments. (B) Western blotting detection of PnpA in WT and ΔpnpA strains (upper panel; ECPs and cells equivalent to 0.3 ml of culture). rPnpA (0.2 µg) was used as control. Lower panel, total protein loading (Ponceau S). Blot shown is representative of three independent experiments. (C) Deletion of pnpA does not affect bacterial growth. Phdp MT1415 and MT1415ΔpnpA strains were grown in TSB-1 at 25°C. Growth curves were generated from three replicates for each strain. Result shown is representative of two independent experiments. (D) Total ion current (TIC) of digested and reduced PG from wild type Phdp MT1415 (top) and MT1415ΔpnpA (middle and bottom; (1) and (2) correspond to two independent cultures of Phdp MT1415ΔpnpA). (E) Deletion of pnpA does not affect bacterial morphology. Bacteria labeled with WGA-Alexa488 (upper panel; scale bar 2 µm). The lenght of at least 150 bacteria from 2 independent experiments was measured and graphed (lower panel, mean lenght ± SD). Statistical significance was tested by Student’s t-test and no differences were observed. (F) Total ion current (TIC) of digested and reduced PG of Phdp previously incubated with Vehicle, PnpA or catalytically inactive PnpAC324A; correspondent reduced supernatants are shown in Fig. S5.

### PnpA has hydrolytic activity towards *Vibrio anguillarum* and *Vibrio vulnificus* PG

The fact that PnpA is actively secreted into the extracellular medium and has no enzymatic activity for *Phdp* PG, raised the possibility that it could cleave PG from other bacteria, functioning as a virulence factor against competing bacteria or as part of a mechanism to acquire nutrients, e.g., muropeptides from dead bacteria. To address this issue, whole sacculi from several Gram-positive or Gram-negative bacteria were isolated and incubated *in vitro* with recombinant PnpA or catalytically inactive PnpA (PnpA^C324A^) (Figs. 7 and S4B-J). Interestingly, only sacculi from *V. anguillarum* and *V. vulnificus* were sensitive to the action of PnpA (Figs. 7, S4B-C and S5). Analysis of the insoluble sacculi resulting from digestion with PnpA showed the appearance of novel muropeptides, not present after incubation with inactive PnpA^C324A^ or vehicle (Fig. 7; Table 2). *V. anguillarum* and *V. vulnificus* PG present a very simple muropeptide composition with three major muropeptides, the monomer GM-tetrapeptide (GM4), the dimer GM4-GM4 and the anhydro-dimer (GM4-GanhM4 and GanhM4-GM4). The high proportion of anhydro-muropeptides indicates that *V. vulnificus* has a PG with short glycan chains (Table 2). PnpA treatment led to the appearance of 4 new muropeptides GM2, GanhM2, GM4-mDapA and GanhM4-mDapA. GM2 and GanhM2 products are consistent with the hydrolysis of the γ-D-glutamyl-*meso*-diaminopimelic acid bond. The presence of GM4-mDapA and GanhM4-mDapA are also consistent with the hydrolysis of a GM4-GM4 or a GM4-GanhM4 dimer at the γ-D-glutamyl-*meso*-diaminopimelic acid bond at one of the two stem peptides.

**FIG 7.**
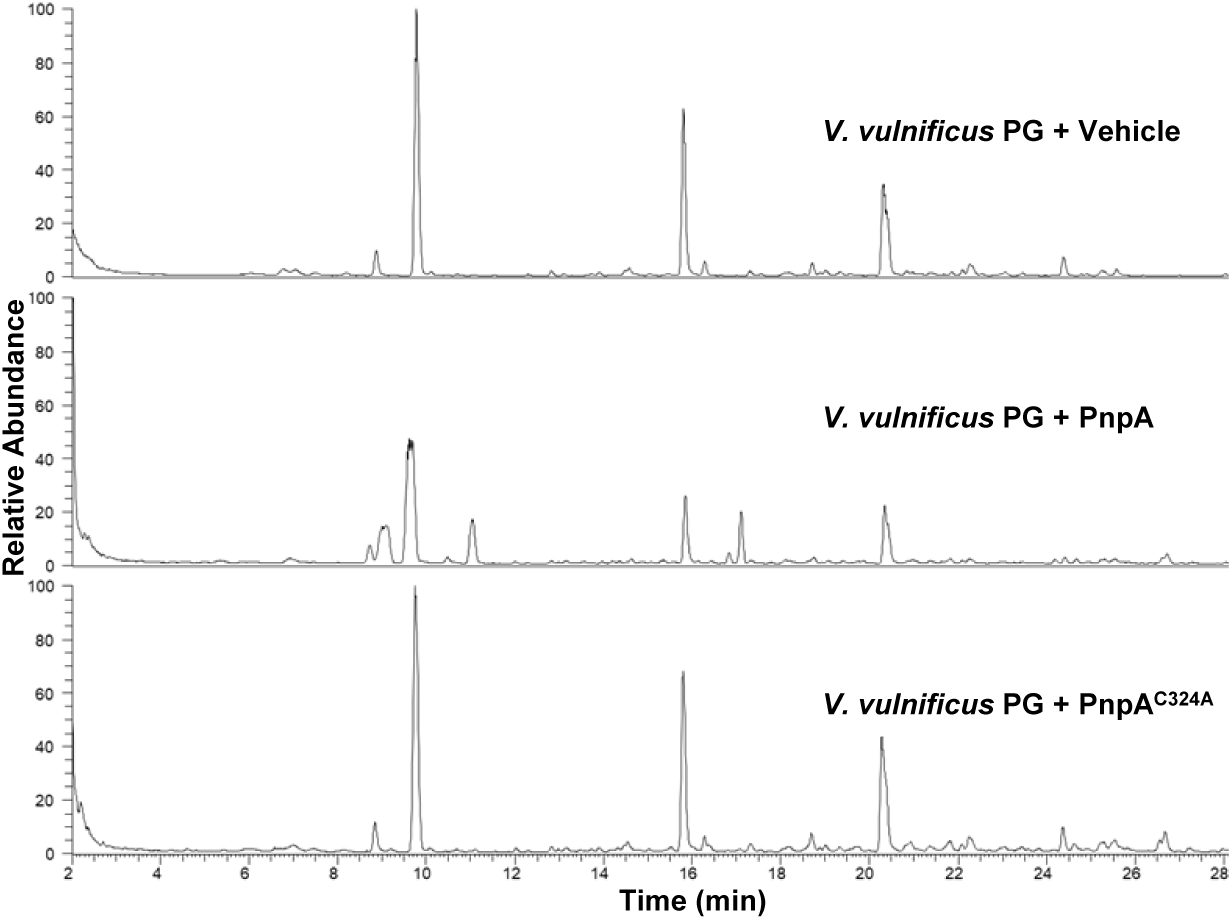
Total ion current (TIC) of digested and reduced PG of *V. vulnificus* previously incubated with Vehicle, PnpA or catalytically inactive PnpA^C324A^; correspondent reduced supernatants are shown in Fig. S5.

Analysis of the products released from the *V. vulnificus* PG, identified two main tetrasaccharides substituted with the L-alanine-D-glutamate dipeptide (GM2-GanhM2) and/or a remain of the dimer cross link (GM4-GanhM4-mDapA; Figs. 7 and S5; Table 3). Additionally, the GanhM2 monomer, the remains of the monomer stem peptide mDapA and of dimer cross-link mDapA-mDapA were also released confirming that PnpA is indeed a γ-D-glutamyl-*meso*-diaminopimeate endopeptidase (Figs. 7 and S5; Table 3).

**TABLE 3.**
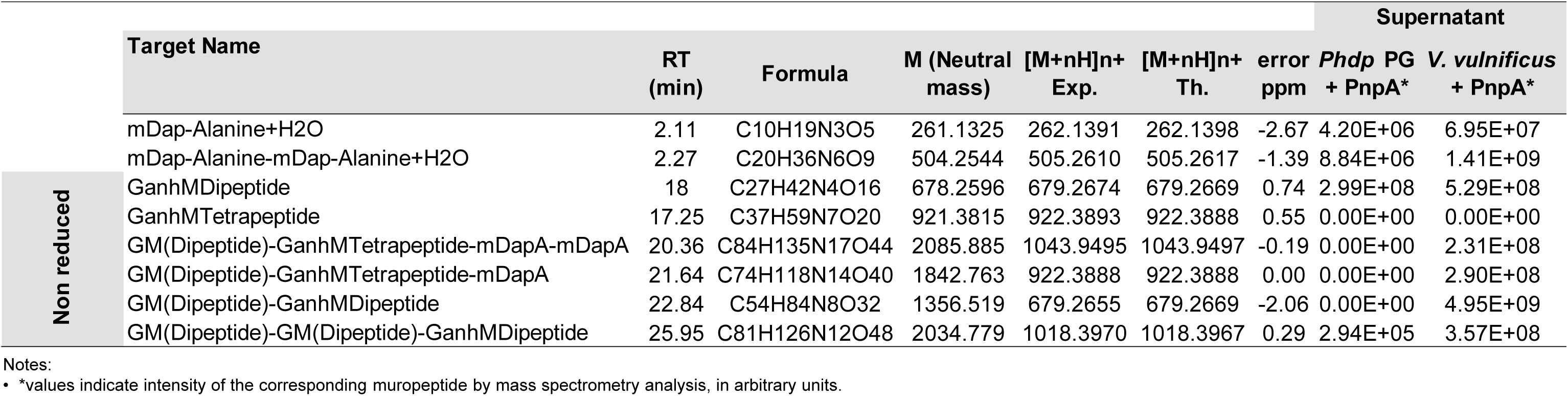
Analysis of PG reduced supernatants from *Phdp* and *Vibrio vulnificus* after incubation with PnpA

In order to assess whether PnpA could inhibit the growth of competitor bacteria, the growth of *V. vulnificus* was monitored in the presence of PnpA (5ug ml^-1^), and no growth inhibition was observed (Fig. S5A). To test the hypothesis that an additional factor secreted by *Phdp* could assist PnpA in reaching the PG, the growth of *V. vulnificus* was monitored in the presence of ECPs from wild type or Δ*pnpA Phdp* (Fig. S5B) and in co-culture experiments (Fig. S5C). No growth inhibition was observed in any of these experiments. Finally, it was tested whether PnpA was able to inhibit the growth of *V. vulnificus* in the presence of EDTA, an external membrane-permeabilizing agent used to mimic conditions that may be encountered in the host, and no effect on growth was observed (Fig. S5D).

## DISCUSSION

In this work, the structural and functional characterization of PnpA, an NlpC/P60 family peptidase secreted by *Photobacterium damselae* subsp. *piscicida* (*Phdp*) is reported. PnpA is not involved in *Phdp* cell wall biogenesis and does not cleave *Phdp* PG, but degrades the PG of *V. anguillarum* and *V. vulnificus*, two bacterial species that share the same hosts and/or environment as *Phdp*. Based on these observations, it is proposed that PnpA may allow *Phdp* to fight competitors or to acquire nutrients from dead co-inhabitants.

Many cysteine peptidases containing the NlpC/P60 domain were characterized to date (1, 2, 24, 26, 30, 33-35), several of which displaying a four-domain organization similar to PnpA. However, until now, only the three-dimensional structure of DvLysin from *Desulfovibrio vulgaris* was reported, with a N-terminal “c-clip” or “N_NLPC_P60” stabilizing domain, two SH3b domains and a C-terminal NlpC/p60 cysteine peptidase domain (26). Furthermore, among the known DvLysin and PnpA orthologs, only EcgA from *Salmonella enterica* serovar Thyphimurium was functionally characterized (56). Although the three molecules are very similar (25-27% amino acid sequence identity) (Fig. S7A), DvLysin does not have the insertion found in PnpA and EcgA and that in PnpA closes the side of the catalytic groove opposed to the RT loop (Figs. 2, 3 and S7B). Despite these differences, residues involved in substrate binding in DvLysin (26) are conserved in PnpA and EcgA (Figs. 3C and S7B), in agreement with their specificity for the γ-D-glutamyl-meso-diaminopimelic acid bond (Fig. 5 and (26, 56)). However, unlike DvLysin (26) and EcgA (56), which were more active towards tetra- and tri-muropeptides, respectively, PnpA showed activity towards penta-, tetra- and tri-peptides (Fig. 5).

So far, the cellular localization of DvLysin and its function in *D. vulgaris* cell wall biogenesis remains unknown (26). Regarding EcgA, its expression is induced when *S*. Typhimurium is inside eukaryotic cells, localizing in the inner and outer membranes where it plays a role in PG remodeling and contributes to *S*. Typhimurium virulence (56). In contrast, PnpA is secreted by the T2SS into the extracellular medium (Fig. 4) and deletion of *pnpA* does not affect *Phdp* growth, PG composition and morphology (Fig. 6C-E). Accordingly, PnpA has no *in vitro* hydrolytic activity towards *Phdp* sacculi (Figs. 6F and S4A). Altogether, these results suggest that PnpA is not involved in *Phdp* cell wall biogenesis.

The resistance of *Phdp* PG to the activity of PnpA is in sharp contrast with the ability of PnpA to hydrolyze penta-, tetra- and tri-muropeptides, since the chemical composition of *Phdp* PG suggested that it would be a target of PnpA. This unexpected resistance to PnpA was not exclusively observed with PG from *Phdp*, as it also occurred when using sacculi from multiple bacterial species (Fig. S4). In fact, PG from *V. anguillarum* and *V. vulnificus* were sensitive to the activity of PnpA, despite having a PG composition characteristic of Gram-negative bacteria and similar to the composition of some PG shown to be resistant to PnpA hydrolysis. Hence, PnpA specificity for *V. anguillarum* and *V. vulnificus* PG cannot be explained by their muropeptide composition and may be related to specific three-dimensional features of the PG mesh. Accordingly, the analysis of the *V. anguillarum* and *V. vulnificus* PG composition shows that these two species have a high proportion of anhydro-muropeptides, a trademark of the end of glycans, indicating that their glycan chains are rather short compared to other Gram-negative bacteria. Consequently, structural analysis of the products released upon incubation of the sacculi of *V. anguillarum* and *V. vulnificus* with PnpA, identified a high proportion of the tetrasaccharide GM2-GanhM2. This suggests that the PG of *V. anguillarum* and *V. vulnificus* is enriched in tetrasaccharides. The simultaneous release of mDapA-mDapA suggests that these tetrasaccharides are linked to the rest of the PG by one or even two cross-links. These results combined with the rather simple muropeptide composition of *V. anguillarum* and *V. vulnificus* suggests that the vulnerability of *V. anguillarum* and *V. vulnificus* to PnpA might come from the fact that their PGs rely on a high degree of cross-linking of very short glycans. Hence, hydrolysis of the stem peptides by PnpA leads to a rapid destruction of the PG layer while in other Gram-negative species, because they have much longer glycans, PG integrity can be maintained by multiple dimers along the same glycan chain (Fig. 8).

**FIG 8.**
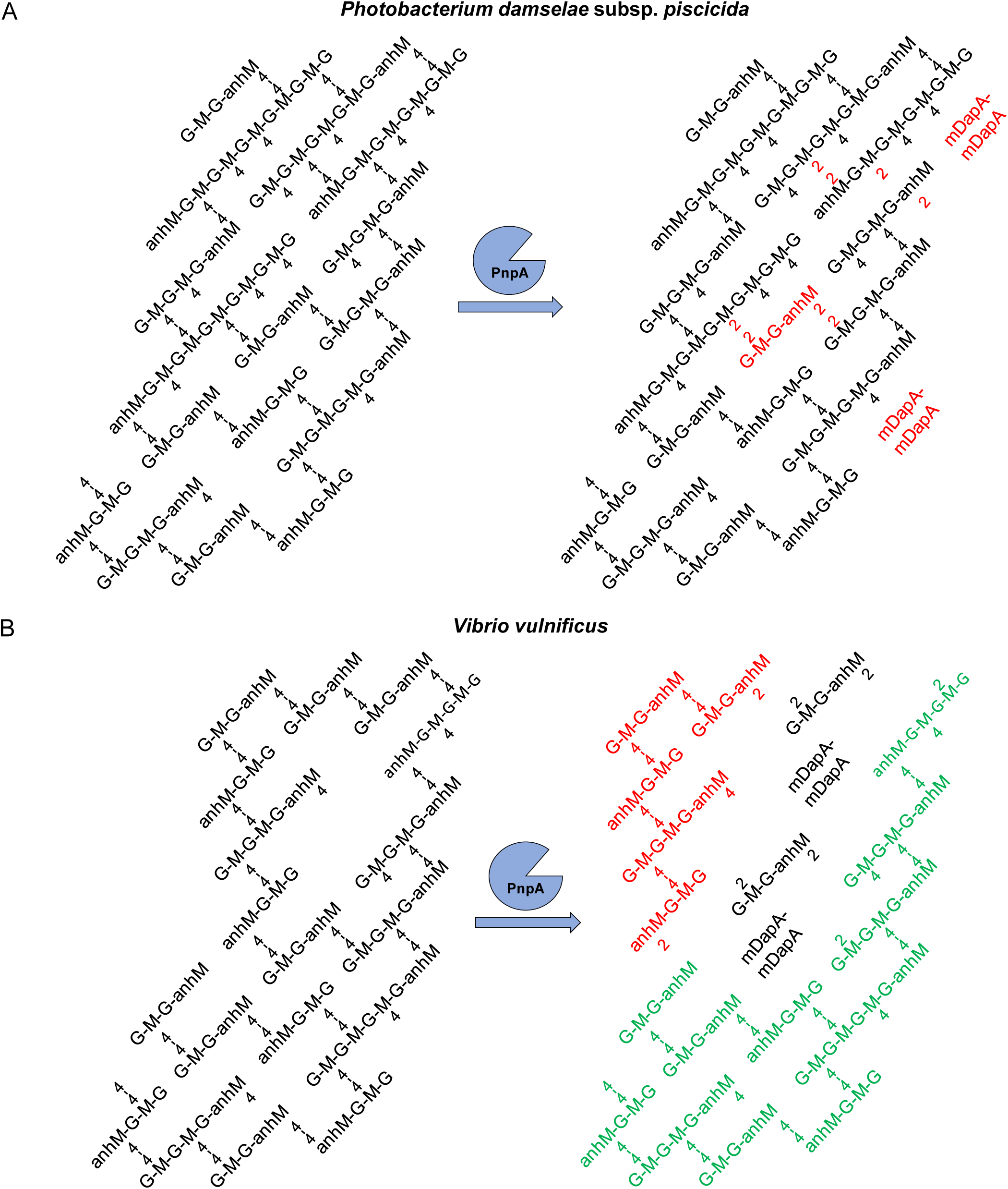
Structural model of Phdp (A) and V. vulnificus (B) PG suggesting that the higher degree of cross-linking of very short glycans in the V. vulnificus PG may explain its vulnerability to the enzymatic activity of PnpA.

Expression levels of *pnpA* in standard culture conditions do not vary between the logarithmic and stationary growth phases (Fig. 1B), but increase under iron-limited conditions or in response to oxidative stress (57). However, *in vivo*, no changes in *pnpA* expression were detected after intraperitoneal infection of sole (*Solea senegalensis*) with *Phdp* (57) and deletion of *pnpA* did not affect *Phdp* virulence in a sea bass (*Dicentrarchus labrax*) intraperitoneal infection model (Fig. S8). This suggests that PnpA is likely dispensable at late systemic phases of *Phdp* infection, but does not exclude a role of PnpA in earlier stages of the infection. It is known that, during the systemic phase of *Phdp*-induced disease, the exotoxin AIP56 plays a major role by neutralizing host phagocytic defenses (43-45, 47, 58). However, little is known about the early stages of the infection. Here, it is shown that PnpA specifically hydrolyses the sacculi of *V. anguillarum* and *V. vulnificus* (Figs. 7, S4B-C and S6), other two enterobacteria present in the marine environment (59-61) and, at least in the case of *V. anguillarum*, reported as infecting the same hosts as *Phdp* (37, 61). This suggests that before reaching the systemic phase, *Phdp* may secrete PnpA to gain competitive growth advantage over bacteria sharing a complex community environment, such as the gastrointestinal tract, or to obtain nutrients, in an environment where nutrient scarcity can compromise its survival, either inside the host or in the water or sediment (40). These strategies have been first described for Gram-positive bacteria (21, 23), which have their PG exposed on the cell surface, accessible to secreted PG hydrolases (10, 11, 22). Gram-negative bacteria, despite having their PG protected by the outer membrane, can inject PG hydrolases, including NlpC/P60 family peptidases, into the periplasm of neighboring bacteria through Type 6 secretion systems (10, 17, 19, 20). The examples of using bacterial exo-hydrolases to target Gram-negative competitors are restricted to predatory bacteria such as myxobacteria (11) and *Bdellovibrio bacteriovorus* (18). Another example where PG hydrolases are secreted to eliminate competing bacteria is that reported for the urogenital pathogenic protozoan *Trichomonas vaginalis* (16), which has acquired by lateral genetic transfer two genes of bacterial origin encoding NlpC/P60 endopeptidases that the parasite secretes to degrade bacterial PG and thus outcompete bacteria from mixed cultures (16). However, it remains unclear how these exo-hydrolases reach the PG of the Gram-negative targets. Here, it was also not clarified how PnpA reaches the PG in *V. vulnificus* and *V. anguillarum* cell wall since no growth inhibition was detected in several *in vitro* tests with *V. vulnificus* (Fig. S5), suggesting that the access of PnpA to the periplasm of competing bacteria may depend on conditions present at specific stages of the *Phdp* life cycle, when *Phdp* and competitors meet together.

## MATERIALS AND METHODS

### Bacterial strains and culture conditions

*Photobacterium damselae* subsp. *piscicida* (*Phdp*) virulent strain MT1415 isolated from sea bass in Italy (45) was cultured at 25°C in tryptic soy broth (TSB) or tryptic soy agar (TSA) supplemented with NaCl to a final concentration of 1% (w/v) (TSB-1 and TSA-1, respectively). The Δ*epsL* and Δ*pnpA* strains were cultured in the same conditions as the wild type. Δ*epsL*+ pEpsL and Δ*pnpA* + pPnpA complemented strains were cultured in TSB-1 and TSA-1 supplemented with 10 µg mL^-1^ of gentamicin (TSB-1_Gm_ and TSA-1_Gm,_ respectively). Stocks of bacteria were maintained at –80°C in TSB-1 supplemented with 15% (v/v) glycerol. To obtain growth curves, bacteria grown on agar plates for 48 h were suspended in TSB-1 or TSB-1_Gm_ at an OD_600_ of 0.5-0.6. These suspensions were inoculated in 20 ml TSB-1 (1:100 dilution). One ml aliquots were removed (in triplicate), transferred to 24-well culture plate and the OD_600_ determined kinetically (1 point/h) using a BioTek Synergy 2 spectrofluorometer (BioTeK U.S., Winooski, VT, USA) at 25°C with continuous slow agitation, for 60-70 h. Growth curves were constructed using GraphPad Prism software (La Jolla, CA, USA).

### Construction of Δ*pnpA* strain

An in-frame (nonpolar) deletion of the almost complete *pnpA* coding sequence was constructed following an allelic exchange procedure as previously described (62). In brief, the 3’ and 5’ flanking sequences were PCR-amplified using suitable primers [Mut_NlpC_1Eco (5’-GC<u>GAATTC</u>GTTTCGATGCGCTGATTAAT-’3); Mut_NlpC_2Bam (5’-GC<u>GGATCC</u>AGCAAAACATCAACAAGTCA-’3); Mut_NlpC_3Bam (5’-GC<u>GGATCC</u>ATAGTTGGTTAATAATGCTA-’3); Mut_NlpC_4Xba (5’-GC<u>TCTAGA</u>TCACGATGGAATAGATAACT-’3) (restriction sites are underlined)]. The PCR products were ligated to obtain an in-frame deletion of ca. 90% of the PnpA coding sequence. The deleted allele construction was cloned into the suicide vector pNidKan containing the *sacB* gene, which confers sucrose sensitivity, and R6K *ori*, which requires the *pir* gene product for replication. The plasmid containing the deleted allele was transferred from *Escherichia coli* S17-1-λ*pir* into the rifampicin-resistant derivative of *Phdp* MT1415 by drop mating for 24 h on TSA plates prepared with seawater. Cells were then scrapped off the plate and selected on TSA supplemented with kanamycin (50 µg mL^-1^) for plasmid integration. A selected Kan^R^ clone was further selected for sucrose resistance (15% [w/v]) for a second recombination event. This led to *Phdp* Δ*pnpA* mutant strain, which was tested by PCR to verify the correct allelic exchange.

### Bacterial cell extracts and extracellular products (ECPs)

*Phdp* were grown in TSB-1 at 25°C with shaking (160 rpm), centrifuged (6000 g, 5 min, 4°C) and the pellets (total cell extracts) and culture supernatants collected. Supernatants were filtered (0.22 µm) to obtain ECPs. For SDS-PAGE, proteins in the ECPs were precipitated with TCA, as previously described (45).

### PnpA Identification

ECPs from *Phdp* strain MT1415 were subjected to SDS-PAGE followed by Coomassie staining. A protein band with approximately 55 kDa was analyzed by MALDI-TOF-MS in a 4800 Proteomics Analyzer (Applied Biosystems) at TOPLAB GmbH. The MS data were used for a Mascot search against the NCBInr sequence database.

### Draft genome sequence of *Phdp* MT1415 and genomic context of *pnpA* locus

To delete the PnpA-encoding gene in *Phdp* MT1415, it was necessary to obtain at least 2 kb of upstream and downstream sequences free of repetitive insertion sequence elements that would compromise the specific recombination steps during allelic exchange. Therefore, the draft genome sequence of strain MT1415 was obtained, using an Illumina platform as previously described (48) and deposited in the GenBank database under accession number SUMH01000000. A comparative analysis was conducted by retrieving the genomic contexts of *pnpA* genes in different *Phdp* and *Phdd* isolates whose draft or complete genomes are available in the GenBank database. The GenBank locus tag numbers of the *pnpA* homologues used in this analysis are: VDA_000779 (*Phdd* type strain CIP 102761), PDPUS_2_00834 (*Phdp* 91-197), PDPJ_2_00460 (*Phdp* OT-51443), BEI67_17705 (*Phdp* L091106-03H), and BDMQ01000002 (*Phdp* DI21). For the *pnpA* negative *Phdd* strain RM-71, the draft genome sequence as source of homologous flanking DNA sequences was used (Accession number NZ_LYBT00000000.1). The DNA sequences were handled with Vector NTI 10.3.0 sequence editor (Invitrogen).

### Recombinant PnpA

*pnpA* ORF (GenBank accession number TJZ86030.1) was amplified from *Phdp* MT1415 genomic DNA using Pfu DNA polymerase (Thermo Scentific) and primers 5’-cgcccATGGATATAAATAAACATTTAATGC-’3 and 5’-gcgctcgagTTTTTCAAATAGATATTTTTC-’3 (target sequences are in uppercase letters) and cloned into pET28a(+) using the NcoI and XhoI restriction sites, in frame with a C-terminal 6xHis-tag. Mutation of Cys324 to alanine was achieved by site directed mutagenesis by inverse PCR using Q5 high fidelity DNA polymerase (New England, BioLabs), pET28-PnpA as template and primers (5’-GCCTCTGGTTTATTAAAAAGGTTATTCAGC-’3 and 5’-ATCATTATTGAAATCCATTCCCCC-’3). Proteins were expressed in *E. coli* BL21 (DE3) CodonPlus-RIL (Stratagene). Four liters of LB medium with 50 μg mL^-1^ kanamycin and 25 μg mL^-1^ chloramphenicol were inoculated with pET28-PnpA or pET28-PnpA^C324A^ transformed bacteria and incubated at 37°C until an OD_600_ of 0.6-0.8. Cultures were cooled at 17°C for 30 min followed by addition of 0.5 mM IPTG to induce protein expression. After 20 h, cells were harvested by centrifugation, resuspended in 50 mM Bis-Tris pH 6.5, 500 mM NaCl and sonicated. Lysates were centrifuged (34957 g, 30 min, 4°C) and the soluble fraction applied to a Ni-NTA column (ABT), followed by anion-exchange chromatography (Bio-Scale Mini Macro-Prep High Q; BioRad). Fractions containing the recombinant proteins were pooled and injected into a size-exclusion chromatography column (Superose12 10/300 GL; GE Healthcare) equilibrated with 50 mM Bis-Tris pH 6.5, 500 mM NaCl. Fractions containing the desired protein were pooled, concentrated to 6-7 mg mL^-1^, frozen in liquid nitrogen and stored at -80°C. Protein concentration was determined in a NanoDrop® ND-1000 UV-Vis Spectrophotometer (Thermo Fisher Scientific) considering the extinction coefficient and the molecular weight calculated by the ProtParam tool (https://web.expasy.org/protparam/).

### Reverse transcription and quantitative PCR (qRT-PCR)

Total RNA was isolated from exponential (OD_600_ of 0.4) and stationary (OD_600_ of 1.2) cultures of *Phdp* strain MT1415. Bacterial pellets were resuspended in 25 mM Tris buffer supplemented with 20% (w/v) glucose and 0.5 M EDTA (pH 8.0) and lysed with phenol acid and glass beads, by vortexing (4°C, 20 min). Lysates were centrifuged at 16000 g (4°C, 5 min) and the top liquid phase was collected. RNA was extracted using the TripleXtractor reagent (Grisp) and treated with DNase I (Turbo DNA-free, Ambion) following the manufacturer’s recommendations. RNA purity and integrity was verified by 1% (w/v) agarose gel electrophoresis in an Experion Automated Electrophoresis System (Bio-Rad). One µg of RNA was reverse-transcribed into cDNA (iScript Kit, Bio-Rad). Quantitative real-time PCR was performed in 20 μl reactions containing 1 μl cDNA, 10 μl *iTaq* Universal SYBR Green Supermix (Bio-Rad Laboratories) and 0.25 μM primers (PnpA forward primer 5’-GGATTTGGCTACCTCGTTCA-’3, PnpA reverse primer 5’-CCCACGGAGCATTAAACATT-’3, 16S forward primer 5’-AACTGGCAGGCTAGAGTCTT-’3 and 16S reverse primer 5’-CACAACCTCCAAGTAGACAT-’3), using the following protocol: 1 cycle at 95°C (3 min) and 40 cycles at 95°C (20 s), 51°C (15 s) and 72°C (30 s). For each condition, three biological replicates were analyzed, each of which with three technical replicates. Data were normalized to the expression values of the housekeeping gene (16S rRNA) and analyzed by the comparative threshold (ΔΔCt) method.

### Anti-PnpA antibody

The quail anti-PnpA antibody was produced at HenBiotech (Coimbra, Portugal, Ref #H003) using recombinant PnpA as immunizing antigen. Quail IgYs were purified from an egg-yolk pool (IgY Grade II/PEG).

### SDS-PAGE and Western blotting

Bacterial cell pellets and ECPs were solubilized in loading buffer (50 mM Tris-HCl pH 8.8, 2% (w/v) SDS, 0.05% (w/v) bromophenol blue, 10% (v/v) glycerol, 2 mM EDTA, and 100 mM DTT) and subjected to SDS-PAGE (63). Proteins were stained with Coomassie Blue or transferred onto nitrocellulose membranes. Transfer efficiency and protein loading were controlled by Ponceau S-staining. Membranes were blocked with 5% (w/v) skim milk in Tris-buffered saline (TBS) containing 0.1% (v/v) Tween 20 (TBS-T), incubated with the anti-PnpA quail antibody (1:10000 dilution) in blocking buffer followed by incubation with an anti-chicken alkaline phosphatase conjugated secondary antibody (Sigma, Ref A9171, 1:10000 dilution) and NBT/BCIP development.

### Crystallization

Initial crystallization hits for PnpA were identified by high-throughput screening performed at the HTX Lab of the EMBL Grenoble Outstation (Grenoble, France). Crystallization experiments for refinement of the initial conditions were carried out using the hanging-drop vapor-diffusion method at 20°C. Crystals were obtained by mixing protein solution (6.7 mg mL^-1^ in 50 mM Bis-Tris pH 6.5, 500 mM NaCl) with an equal volume of crystallization solution (100 mM imidazole pH 8.0, 15% (w/v) PEG 8K). Crystals appeared after 24-48 h. The crystals were cryo-protected by sequential transfer into their crystallization condition with increasing concentrations of ethylene glycol (up to 30% (v/v)) and then flash-frozen in liquid nitrogen prior to data collection.

### Data collection, structure solution and refinement

Diffraction data were collected at beamline Proxima-1 of Synchrotron SOLEIL (Saint-Aubin, France) (64) on a Dectris Pilatus 6M detector (750 images, 0.2° rotation, 0.2 s exposure) and indexed and integrated with XDS (65). Space-group determination, data scaling and merging were performed with POINTLESS and AIMLESS from the CCP4 program suite (66). The structure of PnpA was solved by molecular replacement with Phaser MR as implemented in the CCP4 program suite (66, 67), using the coordinates of a putative gamma-D-glutamyl-L-diamino acid endopeptidase from *Desulfovibrio vugaris* Hildenborough (DvLysin, PDB entry 3m1u, 26% sequence identity) as search model. Phase refinement and initial model building were performed using ARP/wARP (68). Model completion and refinement were done iteratively with COOT (69) and Phenix.refine (70, 71), respectively Refinement and structure-validation statistics are summarized in Table S1. All illustrations of macromolecular models were produced with PyMOL (72). The experimental data were deposited with the Structural Biology Data Grid (73) under accession https://doi.org/10.15785/SBGRID/736.

### *In vitro* muropeptides cleavage assays

To investigate the PnpA enzymatic activity towards PG muropeptides, isolated M3 (GlcNAc-MurNAc-L-Ala-D-Glu-dap), M4 (GlcNAc-MurNAc-L-Ala-D-Glu-dap-D-Ala) and M5 (GlcNAc-MurNAc-L-Ala-D-Glu-dap-D-Ala-D-Ala) muropeptides from *Salmonella enterica* were incubated with 50 µg of PnpA in 50 mM Tris pH 8.0, 300 mM NaCl for 3 h at 37°C. The products of the reaction were analyzed by reverse phase HPLC (Waters 1525 system) as previously described (56).

### Peptidoglycan (PG) purification

Bacteria were grown in TSB-1 at 25°C with shaking (160 rpm) to exponential (OD_600_ of 0.4 - 0.5) or stationary (OD_600_ of 1.2 – 1.4) phases. Bacterial cells (∼10^11^) were centrifuged (4200 g, 10 min, RT), washed twice and resuspended in PBS and immediately mixed 1:1 (v/v) with a boiling solution of 8% SDS, drop by drop. Boiling was maintained for 8 h with stirring, followed by overnight incubation at RT. Samples were centrifuged (150000 g, 40 min, 4°C), the pellets washed three times with ultrapure water (150000 g, 40 min, 4°C), resuspended in 10 mM Tris pH 7.6, 0.06% (w/v) NaCl with or without 100 μg mL^-1^ α-amylase and incubated at 37°C for 90 min. Samples were treated for 2 h at 60°C with 100 μg mL^-1^ Pronase E pre-activated by incubation in the same buffer for 60 min at 60°C. Pronase E digestion was stopped by adding SDS (5.3% (w/v) final concentration) and heating at 100°C for 20 min. PG was recovered by centrifugation (300000 g, 10 min) and washed with ultrapure water.

### Analysis of *Phdp* PG composition and PG cleavage assays

To analyze the PG composition of the *Phdp* MT1415 and MT1415Δ*pnpA* strains, PGs were purified as described above, digested overnight at 37°C in sodium phosphate buffer supplemented with 100 UI of mutanolysin from *Streptomyces globisporus* (ATCC 21553, Sigma) and reduced with NaH_4_B.

After 30 min at RT and centrifugation, the reduced muropeptides were diluted in acidified water with formic acid (FA) and analyzed by HPLC or HPLC/HRMS. HPLC/HRMS was performed on an Ultimate 3000 UHLPC system coupled to a quadrupole orbitrap mass spectrometer (qExactive Focus, Thermo Fisher Scientific). Reduced muropeptides were eluted on an C18 analytical column (Hypersil gold aQ; 1.9 µm, 2.1×150 mm) held at 50°C under a 200 µL min^-1^ flow rate. A binary solvent system composed of acidified water (H_2_O + 0.1% FA; mobile phase A) and acidified acetonitrile (CH_3_CN + 0.1% FA, mobile phase B) was used for chromatographic separation. The composition was linearly increased to 12.5% B over 25 min, increased to 20% B for 5 min and held for an additional 5 min. It was then stepped down to 0% over and held for 10 min to return initial conditions.

qExactive Focus was operated under electrospray ionization in positive mode and data-dependent acquisition mode (ddMS2) control by Xcalibur 4.0. For structural confirmation of muropeptides, HCD fragmentation was set up with a normalized collision energy at 20%. Data were processed both with the software TraceFinder 3.3 (Thermo Fisher Scientific) and Xcalibur 4.0 for peak areas determination.

For testing PnpA activity against macromolecular PG, PG from *Phdp* and several bacteria species, purified as described above, were incubated 100 µg PnpA or inactive PnpA^C324A^ at 37°C overnight in 50 mM Tris pH 8.0, 300 mM NaCl. PG incubated with vehicle were used as controls. After digestion, PG were analyzed by HPLC or HPLC/HRMS, as described above.

### Protein structure accession numbers

The structure factor and atomic coordinate of PnpA are deposited in the RCSB Protein Data Bank (http://www.rcsb.org) with accession number 6SQX.

## Acknowledgements

We are grateful for access to the HTX crystallization facility (Proposal ID: BIOSTRUCTX_8167). The support of the X-ray Crystallography Scientific Platform of i3S (Porto, Portugal) is also acknowledged. This work was financed by FEDER - Fundo Europeu de Desenvolvimento Regional funds through the COMPETE 2020 - Operacional Program for Competitiveness and Internationalization (POCI), Portugal 2020, and by Portuguese funds through FCT - Fundação para a Ciência e a Tecnologia/Ministério da Ciência, Tecnologia e Ensino Superior in the framework of the project POCI-01-0145-FEDER-030018 (PTDC/CVT-CVT/30018/2017). This work had also support from the State Agency for Research (AEI) of Spain co-funded by the FEDER Programme from the European Union (grants AGL2016-79738-R and BIO2016-77639-P) and from the French Government’s Investissement d’Avenir program, Laboratoire d’Excellence “Integrative Biology of Emerging Infectious Diseases” (grant n ANR-10-LABX-62-IBEID; http://www.agence-nationale-recherche.fr/investissements-d-avenir/). AR was support by a post-doctoral fellowship from the Laboratoire d’Excellence “Integrative Biology of Emerging Infectious Diseases and from an Infec-ERA grant (INTRABACWALL - 16-IFEC-0004-03).

Author contributions are as follows: Conceptualization-JL, AdV and NMSdS; Data curation-PJBP; Formal analysis-JL, CP, JA, PJBP, FG-dP, IGB, AdV and NMSdS; Funding acquisition-CRO, FG-dP, IGB, AdV and NMSdS; Investigation-JL, CP, AR, JA, MST, AVB, IR, PJBP, CRO, FG-dP, IGB and AdV; Methodology-JL, CRO, IGB, AdV and NMSdS; Project administration-AdV and NMSdS; Supervision-PJBP, CRO, IGB, AdV and NMSdS; Validation-JL, PJBP, IGB, AdV and NMSdS; Writing (original draft)-JL, CRO, AdV and NMSdS; Writing (review & editing)-JL, CP, JA, PJBP, CRO, FG-dP, IGB, AdV and NMSdS

## FIGURE LEGENDS

**TABLE S1.**
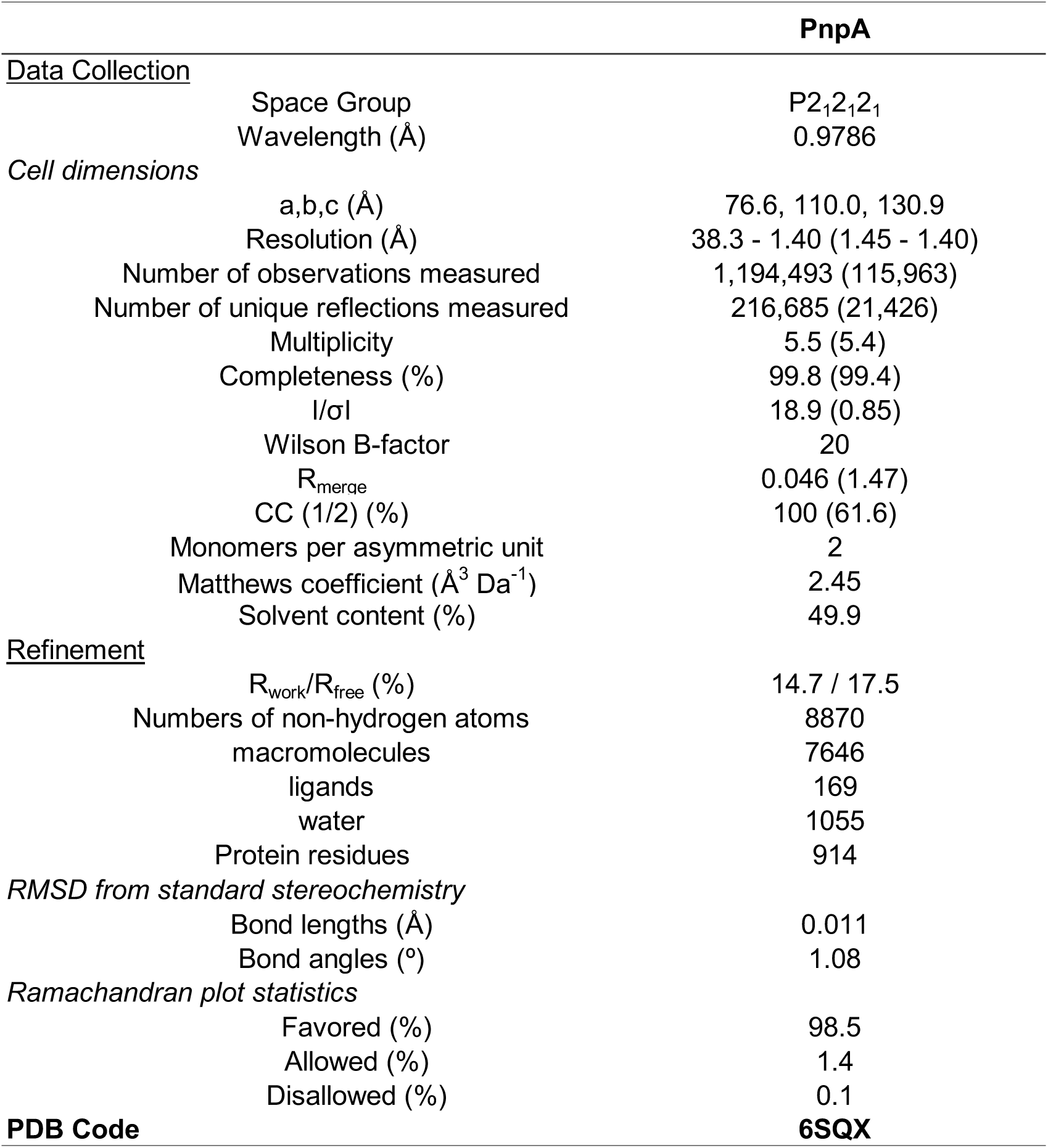
Data collection and refinement statistics of PnpA. The numbers in parentheses are for the highest resolution shell

**TABLE S2.** Structural comparison of PnpA and other bacterial proteins with at least one common domain. Alignments were performed by DALI structural comparison server (Holm, L. Bioinformatics 2019, 35:5326-53-27; Holm, L. In: Gáspári Z. (eds) Structural Bioinformatics. Methods in Molecular Biology, vol 2112. Humana, New York, NY, pp. 29-42) using full-length PnpA as a search probe. For proteins with multiple SH3-like domains or multiple chains, only the best match is shown.

| Aligned substructure(s) | PDB code | Reference | R.m.s.d. (Å) | Aligned length | No. of residues<sup>†</sup> | Z-score<sup>a</sup> | Sequence identity (%) | Comments |
| --- | --- | --- | --- | --- | --- | --- | --- | --- |
| **C-clip + SH3b + NlpC/P60** | 6SQX | This study |  |  |  |  |  | putative gamma-D-glutamyl-L-diamino acid endopeptidase PnpA |
|  | 3M1U | JCSG<sup>‡</sup> | 2.2 | 407 | 420 | 43.3 | 26 | putative gamma-D-glutamyl-L-diamino acid endopeptidase DvLysin |
| **SH3b + NlpC/P60** | 3H41 | Xu et al. (2010) | 2.6 | 251 | 307 | 22.4 | 19 | gamma-D-glutamyl-L-diamino acid endopeptidase Ykfc |
|  | 3NPF | JCSG<sup>‡</sup> | 2.9 | 256 | 305 | 21.2 | 19 | putative dipeptidyl-peptidase VI |
|  | 4R0K | JCSG<sup>‡</sup> | 3 | 253 | 304 | 21.1 | 20 | putative dipeptidyl-peptidase VI |
|  | 3PVQ | JCSG<sup>‡</sup> | 2.8 | 246 | 297 | 20.5 | 20 | putative dipeptidyl-peptidase VI |
|  | 2HBW | Xu et al. (2009) | 2.4 | 182 | 220 | 14 | 19 | putative endopeptidase AvPCP |
|  | 2EVR | Xu et al. (2009) | 2.4 | 182 | 222 | 13.9 | 19 | putative gamma-D-glutamyl-L-diamino acid endopeptidase NpPCP |
|  | 6BIQ | Pinheiro J. et al. (2018) | 4.7 | 159 | 266 | 10.9 | 16 | NlpC/P60 D, L endopeptidase (NlpC_A2) |
|  | 6BIO | Pinheiro J. et al. (2018) | 4.5 | 159 | 278 | 10.7 | 15 | NlpC/P60 D, L endopeptidase (NlpC_A1) |
| **Lysozyme-Like + NlpC/P60** | 4FDY | Xu et al. (2014) | 2.3 | 115 | 295 | 11.5 | 21 | bifunctional Cell Wall Hydrolase CwIT |
|  | 4HPE | JCSG<sup>‡</sup> | 2.7 | 121 | 290 | 11.3 | 22 | putative cell wall hydrolase |
| **Coil-Coil Domain + NlpC/P60** | 6B8C | Kim B. et al. (2019) | 2.3 | 108 | 117 | 10.7 | 23 | NlpC/p60 domain of peptidoglycan hydrolase SagA |
| **LysM + NlpC/P60** | 4XCM | Wong et al. (2015) | 2.7 | 115 | 218 | 11.2 | 19 | putative NlpC/P60 D, L endopeptidase LysM |
| **NlpC/P60** | 3I86 | Unpublished | 2.4 | 117 | 136 | 11.1 | 26 | P60 Domain |
|  | 2K1G | Aramini et al. (2008) | 2.2 | 115 | 129 | 10.9 | 24 | NlpC/P60 domain of lipoprotein Spr |
|  | 3GT2 | Unpublished | 2.3 | 114 | 135 | 10.4 | 21 | P60 Domain |
|  | 4JXB | Both et al. (2014) | 2.5 | 113 | 130 | 10.2 | 22 | RipD, a non-catalytic NlpC/p60 domain protein |
|  | 3PBI | Both et al. (2011) | 2.6 | 126 | 199 | 9.4 | 25 | peptidoglycan hydrolase RipB |
|  | 3NE0 | Ruggiero et al. (2010) | 4.1 | 130 | 208 | 8.4 | 24 | RipA, a Mycobacterial Enzyme |
|  | 2XIV | Unpublished | 4.1 | 129 | 207 | 8.3 | 23 | Rv1477, Hypothetical Invasion Protein |
|  | 3PBC | Both et al. (2011) | 4.2 | 129 | 208 | 8.3 | 25 | Peptidase module of the peptidoglycan hydrolase RipA |
|  | 4Q4N | Squeglia et al. (2014) | 4.5 | 131 | 208 | 8.3 | 22 | RipA, a Mycobacterial Enzyme |
|  | 4EYZ | Levy-Assaraf et al. (2013) | 3.3 | 140 | 246 | 7.8 | 15 | Cellulosome-related protein module |
|  | 6IST | Unpublished | 3.4 | 119 | 213 | 7.5 | 8 | Endolysin LysIME-EF1 |

<sup>†</sup> Number of residues present in the model used for comparison. <sup>‡</sup> Joint Center for Structural Genomics. <sup>§</sup> Northeast Structural Genomics Consortium. <sup>§</sup> Ontario Centre for Structural Proteomics. <sup>a</sup> Z-score is a measure of the quality of the alignment. The higher the Z-score, the more homologous are the structures.

**FIG S1.**
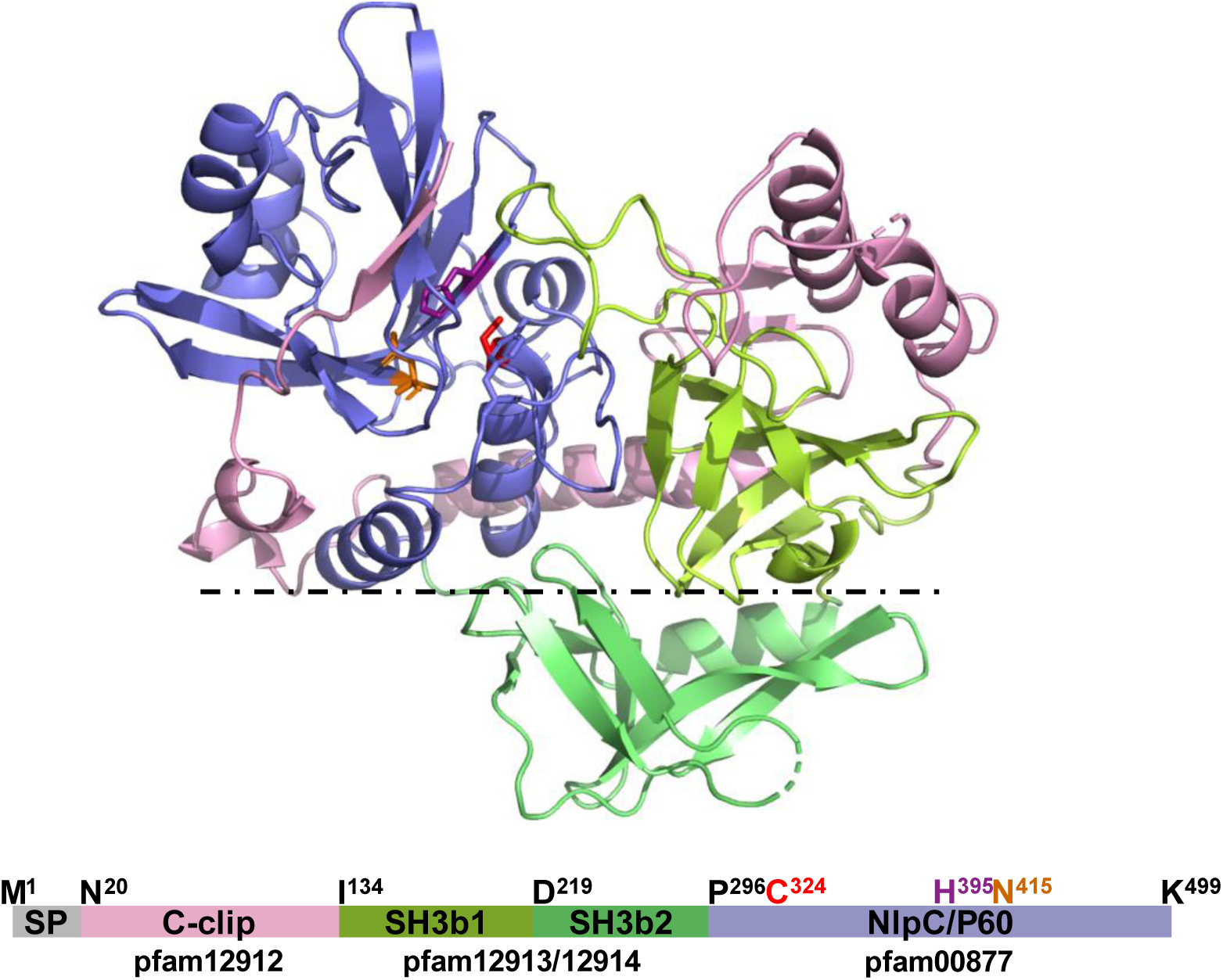
Cartoon representation of the three-dimensional structure of PnpA highlighting the protrusion of the SH3b2 domain relative to the plane defined by the other domains. Color code as in Fig. 2.

**FIG S2.**
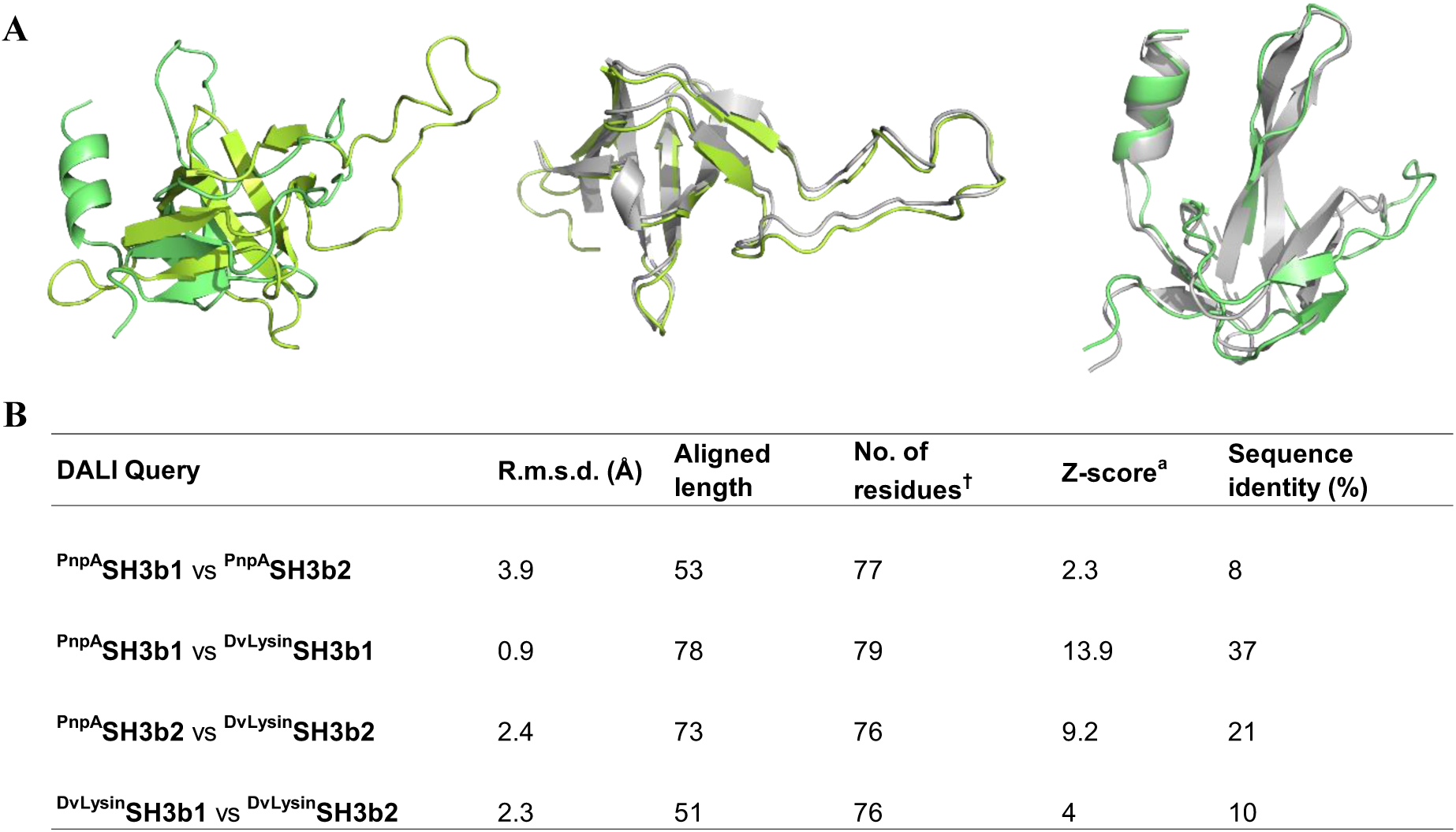
Structural comparison of the SH3b domains of PnpA and DvLysin (PDB entry : 31MU). (A) Superposition of SH3b1 (light green) and SH3b2 (dark green) domains from PnpA (left), of SH3b1 domains from PnpA (light green) and DvLysin (gray) (middle), and of SH3b2 domains from PnpA (dark green) and DvLysin (gray) (right). (B) Results of the pairwise superposition of SH3b domains using the DALI server (Holm, L. Bioinformatics 2019, 35:5326-53-27; Holm, L. In: Gáspári Z. (eds) Structural Bioinformatics. Methods in Molecular Biology, vol 2112. Humana, New York, NY, pp. 29-42). a Z-score is a measure of the quality of the alignment. The higher the Z-score, the more homologous are the structures.

**FIG S3.**
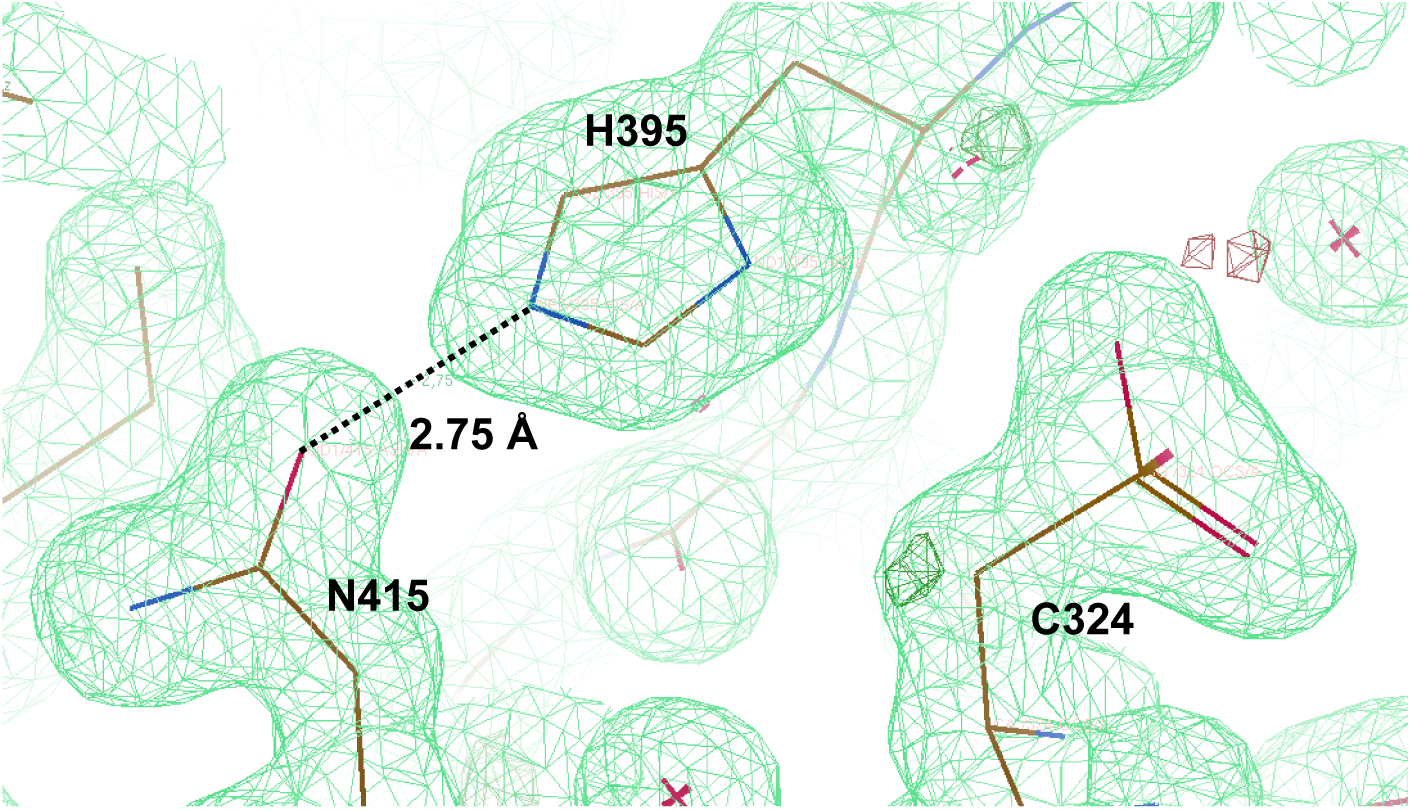
Active site region of Phdp PnpA with superposed 2Fo - Fc electron density map (green mesh) contoured at 1.0σ. The contact normally established between the imidazole ring of H395 and the side chain of N415 is represented by a dashed line and the respective distance (in Å) indicated.

**FIG S4.**
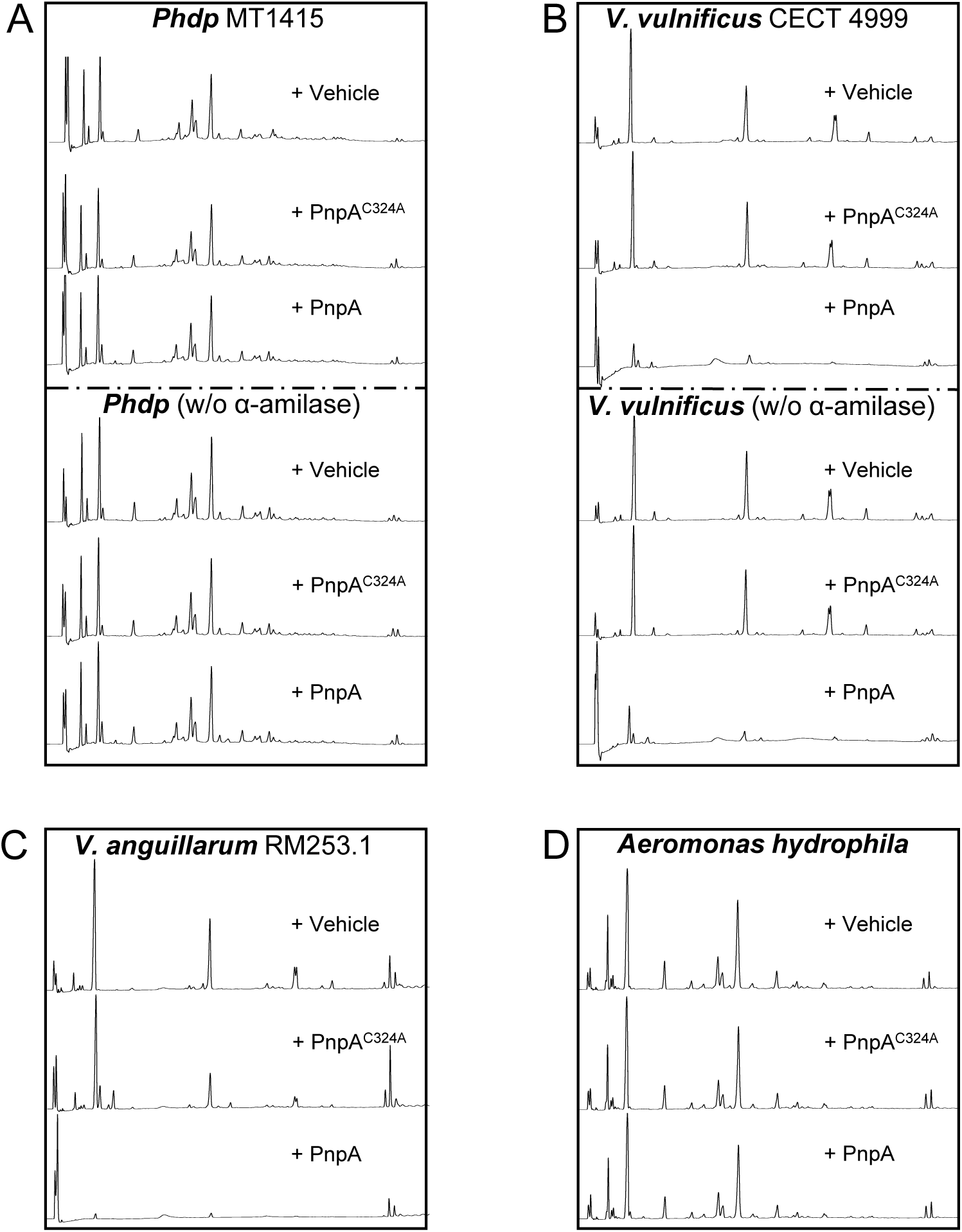
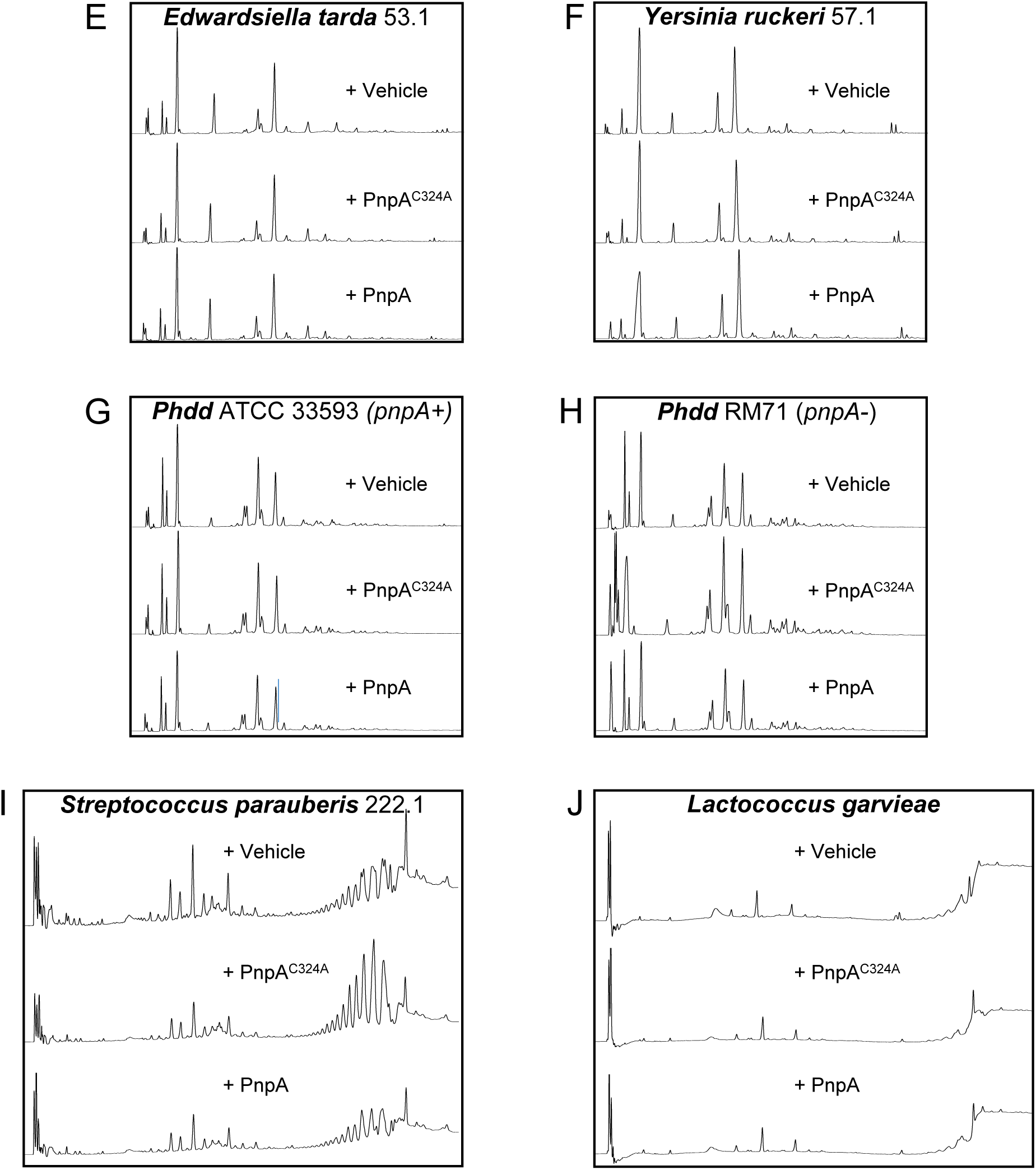
PnpA degrades the peptidoglycan (PG) of Vibrio vulnificus (B) and V. anguillarum (C), but not the PG of Photobacterium damselae subsp. piscicida (A), or of other Gram-negative (D-H) and Gram-positive bacteria (I,J). Unless stated otherwise, incubations were performed after treating the PG with α-amilase. In the case of Phdp and V. vulnificus, digestions of PG not treated with α-amilase were also performed and no differences were obtained.

**FIG S5.**
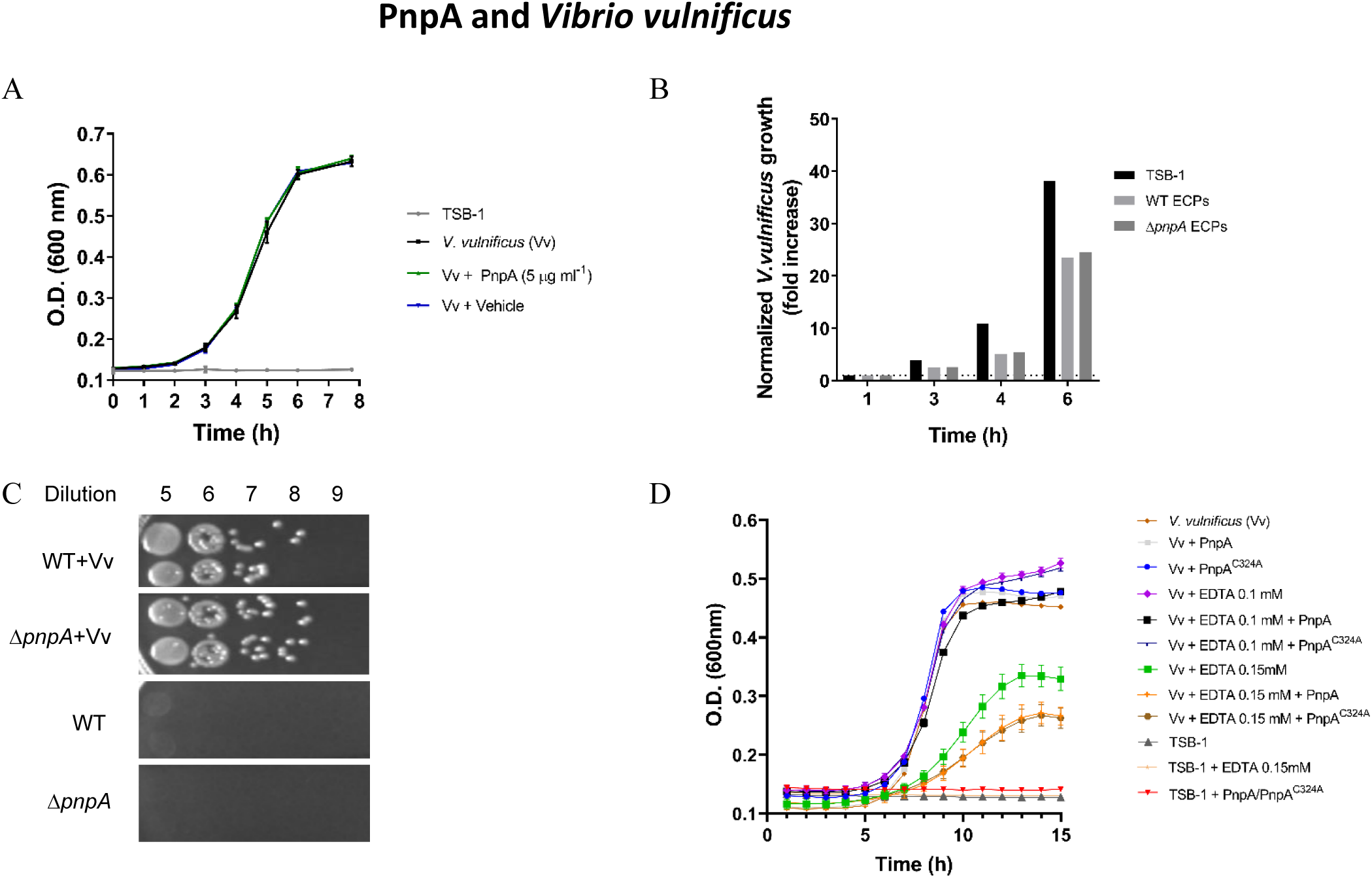
PnpA does not affect Vibrio vulnificus growth. (A) Recombinant PnpA does not inhibit V. vulnificus growth. V. vulnificus strain CECT 4999 (Vv) was grown in TSA-1 for 24 h at 25°C and suspended in TSB-1 at an OD_600_ of 0.5-0.6. This suspension was inoculated in 1 ml TSB-1, TSB-1+5 µg mL^-1^ PnpA or TSB-1+vehicle at 1:50 dilution in wells of a 24-well plate. The plate was incubated at 24°C with shaking and the OD_600_ was read using Synergy plate reader. Growth curves were generated from the three replicates for each condition. Result shown is representative of two independent experiments. (B) PnpA does not affect V. vulnificus growth in the presence of other extracellular products secreted by Phdp. Phdp MT1415 and MT1415ΔpnpA strains were grown in TSA-1 for 48 h and V. vulnificus (Vv, CECT 4999) was grown in the same media for 24 h. MT1415 and MT1415ΔpnpA were suspended in TSB-1 at an OD_600_ of 0.5, inoculated 1:100 in TSB-1 and grown at 25°C until an OD_600_ of 0.8. ECPs were collected by centrifugation and filtered with 0.2 µm filter. Vv were suspended in TSB-1 at an OD_600_ of 0.5 and were inoculated in 20 ml TSB-1 or in ECPs from MT1415 or MT1415ΔpnpA at 1:50 and grown at 25°C. The OD_600_ was measured at the indicated times and normalized for the OD_600_ measured at 1 h. Result shown is representative of two independent experiments. (C) Growth of V. vulnificus is not affected by co-culture with Phdp. Phdp MT1415 and MT1415ΔpnpA strains were grown in TSA-1 plates for 48 h and V. vulnificus (VV, CECT 4999) was grown in the same media for 24 h. MT1415 WT and ΔpnpA were suspended in TSB-1 at an OD_600_ of 0.5, inoculated 1:50 in TSB-1 and grown overnight. The cultures were refreshed in TSB-1 to an OD_600_ of 0.25 and grown until an OD_600_ of 0.5. Vv was suspended in TSB-1 to an OD_600_ of 0.5, inoculated in the WT and ΔpnpA cultures at 1:50 and grown at 25°C for 6 h. Co-cultures were serial diluted (1:10) in TSB-1 and spotted (10 µL dots) in TSA-1 or TCBS agar. Note that TCBS-agar allows the growth of Vv but not of Phdp. Result shown is representative of two independent experiments. (D) PnpA does not affect V. vulnificus growth in the presence of the outer membrane permeabilizer EDTA. V. Vulnificus grown in TSA-1 for 24 h at 25°C and suspended in TSB-1 at an OD_600_ of 0.5-0.6 was inoculated in 1 ml TSB-1, TSB-1+0.10 mM EDTA or TSB-1+0.15 mM EDTA, or the same media containing 30 µg mL^-1^ of PnpA or inactive PnpAC324A at 1:50 dilution, in an 24-well plate. TSB-1 (Blank), TSB-1+0.15 mM EDTA and TSB-1+30 µg mL^-1^ PnpA+30 µg mL^-1^ PnpAC324A were used as controls. The plate was incubated at 25°C with shaking and the OD_600_ was read using Synergy plate reader. Growth curves were generated from three replicates for the conditions containing Vv and from a single read for the controls. Result shown is from one experiment.

**FIG S6.**
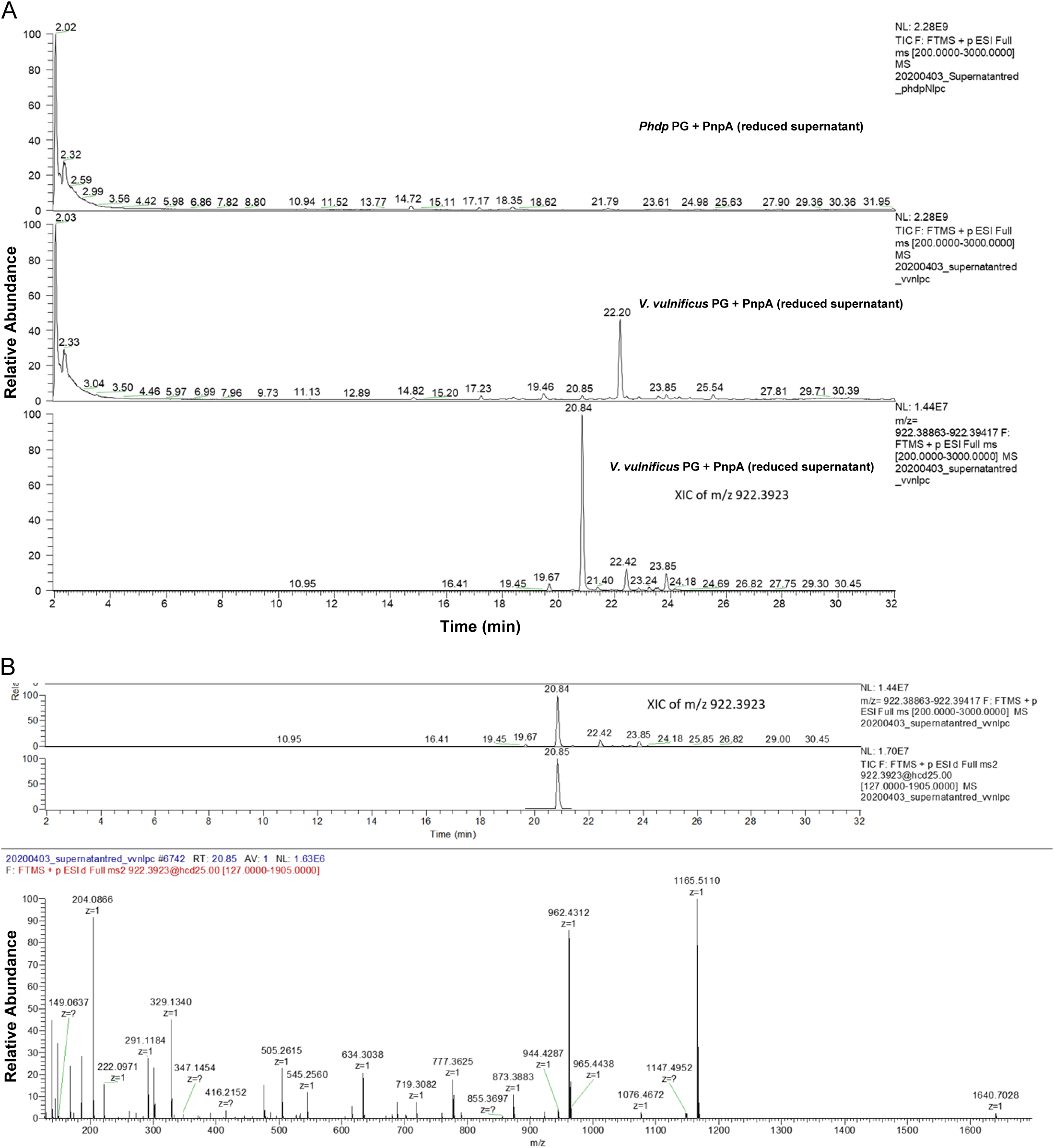
(A) Reduced supernatants from peptidoglycan (PG) of Phdp and Vibrio vulnificus incubated with PnpA. At the bottom, ion extract of m/z 922.3923 in V. vulnificus supernatant; (B) Ion extract of m/z 922.3923 shown in (A) and its MS/MS fragmentation spectrum.

**FIG S7.**
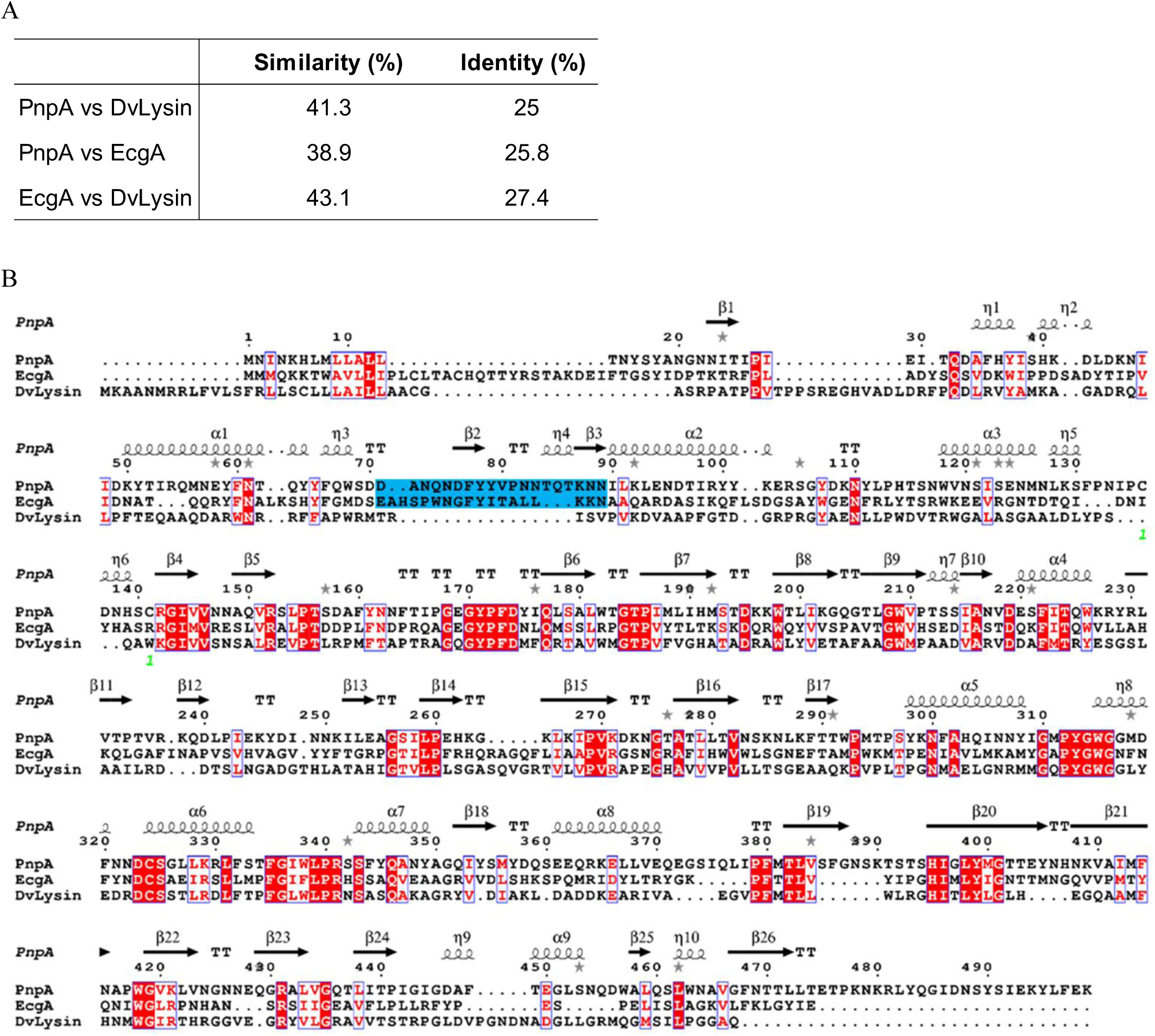
Amino acid similarity and identity determined with EMBOSS (A) and amino acid sequence alignment (B) of PnpA, EcgA from Salmonella enterica serovar Typhimurium and DvLysin from Desulfovibrio vulgaris Hildenborough. Alignment was performed using ClustalW (Thompson, J. D. et al. Nucleic Acids Research 1994, 22(22):4673-4680). Strictly conserved residues are in white on a red background. Partially conserved amino acids are boxed. Secondary structure elements from PnpA are indicated above the alignment. The number (green) below the sequences indicate disulfide bridge present in PnpA structure. The cyan box highlights the antiparallel β-sheet (β2 and β3) and a 310 helix (η4) in the c-clip domain of PnpA, EcgA that is absent in DvLysin. This figure was generated using the ESPript server (ESPript - http://espript.ibcp.fr; Robert, X. and Gouet, P. Nucleic Acids Research 2014, 42:W320-W324).

**FIG S8.**
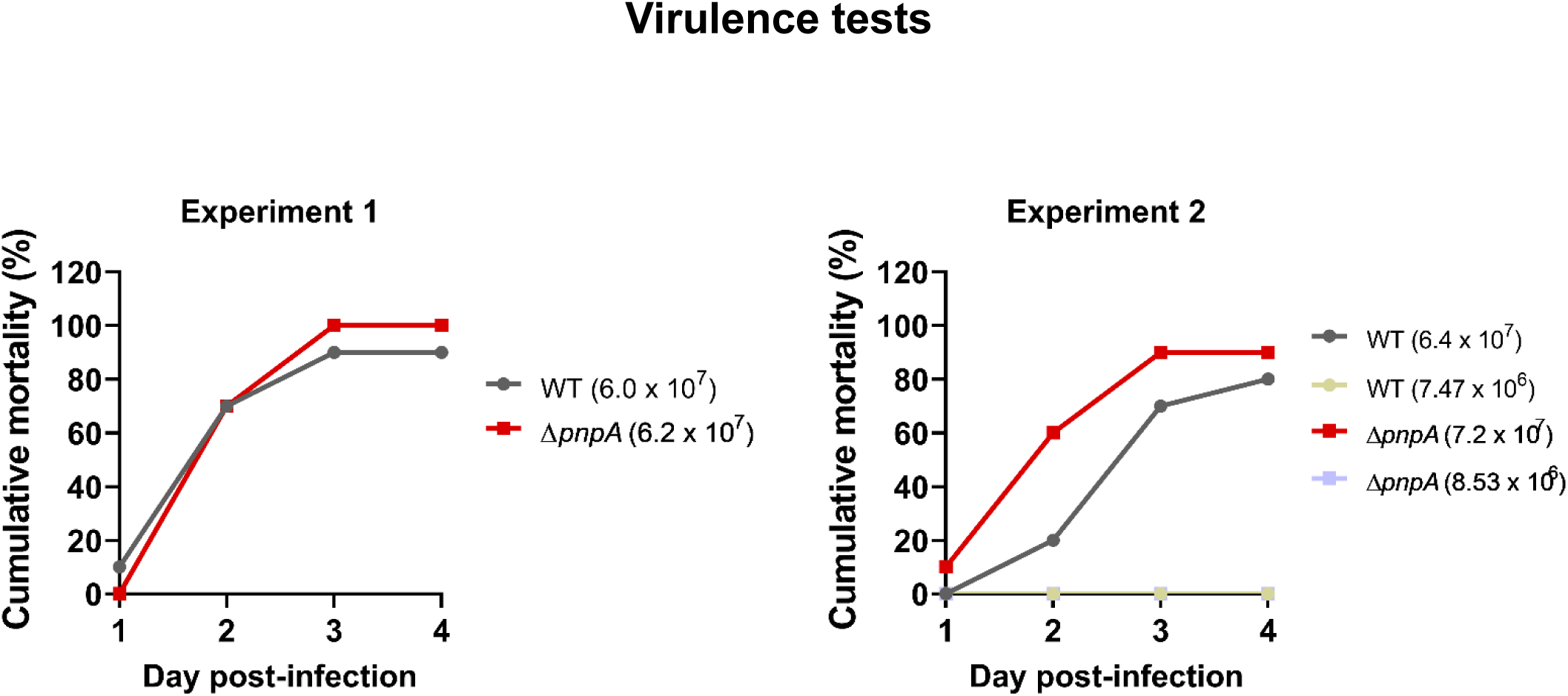
Deletion of pnpA does not affect sea bass mortality after i.p. infection. Cumulative mortality of juvenile sea bass (average weight of 19.63±3.63 g and 20.35±3.86 g for experiments 1 and 2, respectively) injected i.p with the indicated doses of wild type or ΔpnpA strains. The study was carried out in accordance with European and Portuguese legislation for the use of animals for scientific purposes (Directive 2010/63/EU; Decreto-Lei 113/2013). The work was approved by Direcção-Geral de Alimentação e Veterinária (DGAV), the Portuguese authority for animal protection (reference 0421/000/000/2013). Fish were maintained in recirculating aerated sea water at 22°C and previously acclimatized to 23-24°C prior to infection. To prepare the inocula, bacteria grown in TSA-1 plates for 48 h at 25°C were suspended in TSB-1 at an OD_600_ of 0.4 and these suspensions were used to inoculate 100 mL TSB-1 at 1:100 dilution. The cultures were grown for 14-15 h at 25°C with shacking (160 rpm) until reaching an OD_600_ of 0.5-0.6, centrifuged (3220 g, 15 min, 4°C) and resuspended in TSB-1 at an OD_600_ of 0.6. Groups of 10 juvenile sea bass were anesthetized with 0.06% (v/v) ethilenoglycol monophenyl ether and i.p. injected with 100 µl of these suspensions. In experiment 2, 1:10 dilutions of the initial suspensions were also injected. CFUs of each inoculum were determined by plating serial dilution onto TSA-1.

